# Combinatorial transcription factor interactions drive modular gene regulatory networks

**DOI:** 10.64898/2026.05.20.726505

**Authors:** Jialei Duan, Boxun Li, Kartik Kulkarni, Gabriela Orquera-Tornakian, Daryl Barth, Lei Wang, Vrushali Pandit, Joel Liou, Nikhil V Munshi, Gary C Hon

**Author notes:** Equal contribution (Jialei Duan, Boxun Li, Kartik Kulkarni). **Data availability** The single cell RNA sequencing data reported in this paper were deposited in NCBI Gene Expression Omnibus (GEO) under the accession number GSE225838. To review, go to https://www.ncbi.nlm.nih.gov/geo/query/acc.cgi?acc=GSE225838 and enter token unmjwmaibbwxpkd into the box.

## Abstract

Transcription factors (**TFs**) cooperatively drive gene regulatory networks (**GRNs**) to establish transcriptional states. Forced induction of TFs in combination can reprogram cell state by supplanting existing GRNs. Thus, TFs and GRNs are the building blocks to engineering transcriptional state. However, one key challenge is that the relationship between TF combinations and GRNs remains largely uncharacterized and difficult to accurately predict. Here, we apply single-cell overexpression screens to map the combinatorial activities of ∼100 TFs to gene expression states. Our analysis identifies diverse TF combinations driving cell-type specific regulatory programs. Notably, different TF combinations induce shared gene sets with cell-type specific functions, suggesting a modular regulatory architecture of the transcriptome. Furthermore, we define pairwise TF interactions and show that cooperative interactions improve transcriptional reprogramming. Finally, we developed tools to predict combinatorial TF phenotypes. These findings improve our understanding of cell state and how to manipulate it for biomedical applications.

**HIGHLIGHTS:**

- Combinatorial over-expression screens for ∼100 transcription factors (TFs).
- Diverse TF combinations drive cell-type specific regulatory programs.
- TF regulatory networks reveal a modular regulatory architecture of the transcriptome.
- TF-TF interactions and predictive models enhance reprogramming cocktails.

## INTRODUCTION

Single-cell genomics is rapidly defining transcriptional states across a spectrum of cell types and diseases ^1–3^, providing rigorous evidence that transcriptional states define functional cell states. However, despite these extensive maps, the drivers of gene expression states remain less characterized.

Transcription factors (**TFs**) act cooperatively to drive gene regulatory networks (**GRNs**) and establish a cell’s transcriptional state ^4–7^. Although developmental GRNs are hierarchical and often unidirectional ^8,9^, direct reprogramming by TFs can rewire GRNs to ectopically express lineage-specific genes and potentially create alternative cell types. For example, Oct4, Sox2, and Nanog coordinately regulate pluripotency gene expression in embryonic stem cells ^7^, but forced induction of TF combinations can ectopically activate GRNs in other cells ^10,11^. In extreme cases, this results in functional reprogramming, for example in the conversion of fibroblasts to iPS cells with Oct4, Sox2, Klf4, and Myc ^12,13^. Similarly, in cardiac reprogramming, GHMT (Gata4, Hand2, Mef2c, Tbx5) synergistically activate cardiac enhancers and regulatory programs ^11^. Despite these successes, identifying reprogramming TFs remains challenging for several reasons: the difficulty of engineering reporters of successful reprogramming, the lack of candidate reprogramming factors rooted in developmental biology ^12,14^, and the slow, labor intensive, and low throughput nature of traditional approaches. To address these problems, we and others have developed single-cell screens to identify TF cocktails for transcriptional reprogramming ^15–18^. Our screening method was named Reprogram-Seq (1.0), which we continue to improve here ^16^.

While diverse studies rigorously support TF-mediated reprogramming ^14,19–21^, *functional* reprogramming (full cell state conversion) can be inefficient. In contrast, *transcriptional* reprogramming is more readily attainable and represents a prerequisite for functional reprogramming. In turn, transcriptional reprogramming can be distilled into combinations of GRNs that are activated by combinations of TFs. Although individual TFs drive transcription, most reprogramming cocktails consist of 2 or more TFs ^22^ that often interact non-linearly to produce emergent properties ^23,24^. For example, cooperativity between TFs can change the DNA binding specificities of paired TFs ^25^, and specific TF pairs reprogram distinct neuronal subtypes, while individual TFs do not ^26^. These observations indicate that TF pairs represent the basic unit of meaningful GRN reprogramming. Thus, empirical testing of TF cooperativity and understanding these interactions will improve our ability to accurately identify the building blocks necessary for predicting reprogramming cocktails.

While combinatorial TF interactions are important for reprogramming, their nonlinear interactions remain difficult to predict. One common approach to predict *individual* TF activity is to construct GRNs by examining transcriptomic datasets ^27–30^. However, GRN inference remains a difficult computational problem because important parameters are either missing or incomplete, such as the association of TFs, binding motifs, and promoters ^31^. Aside from a handful of successes ^32,33^, the problem of predicting *combinatorial* TF activities is more complex. Despite these known pitfalls, contemporary methods critically rely on predicted GRNs to predict TF cocktails ^29,30,34,35^. Deconvoluting the relationship between TFs and their target genes could be aided by direct perturbation of TF combinations.

Recent studies have comprehensively cataloged the functional activities of ∼1800 human TFs individually ^17,36^, but the space of combinatorial TF activities is too immense to be comprehensively sampled (∼1.6 million combinations of 2 TFs, ∼970 million combinations of 3 TFs). Therefore, systematic cataloging of all possible higher-order combinatorial perturbations is not possible given current technologies. Here, we perform combinatorial perturbation studies of ∼100 TFs in mouse embryonic fibroblasts (**MEFs**) for applications in reprogramming. By measuring the transcriptional outcomes of TF over-expression, these experiments directly measure the gene regulatory networks driven by TFs, define the interactions of TF pairs in the context of reprogramming, and reveal a shared regulatory architecture of common gene modules. An initial study of fitness phenotypes in a loss-of-function paradigm focused analyses on a specific subset of genetic interactions^37^. In the current study, we implement and extend this framework to examine combinatorial interactions between TFs in a gain-of-function, lineage reprogramming model system. Altogether, these approaches represent a generalizable framework for transcriptional, and ultimately functional, reprogramming.

## RESULTS

### Combinatorial over-expression screens for 105 transcription factors

Reprogram-Seq couples combinatorial over-expression of a library of TF open reading frames (ORFs) and single-cell RNA-Seq, for high throughput measurements of TF perturbations and resulting transcriptional states in individual cells (**Figure 1A**) ^16^. We introduced TF-specific barcodes immediately downstream of the TF transgene to minimize TF/barcode recombination during lentivirus production ^38,39^. A PCR handle upstream of the TF barcode facilitates construction of perturbation libraries. We synthesized codon optimized TFs and barcodes, and validated synthesis by sequencing. To reduce the heterogeneity of lentiviral titers across ORFs, we developed an iterative sequencing-based strategy to normalize relative viral titers across pools of TFs, and we incorporated a GFP control to control absolute viral titers (see Methods) ^40–42^. In benchmarking experiments, we confirm the ability to precisely control and evaluate viral titer prior to sequencing (**Figure S1A-D)**.

**Figure 1:**
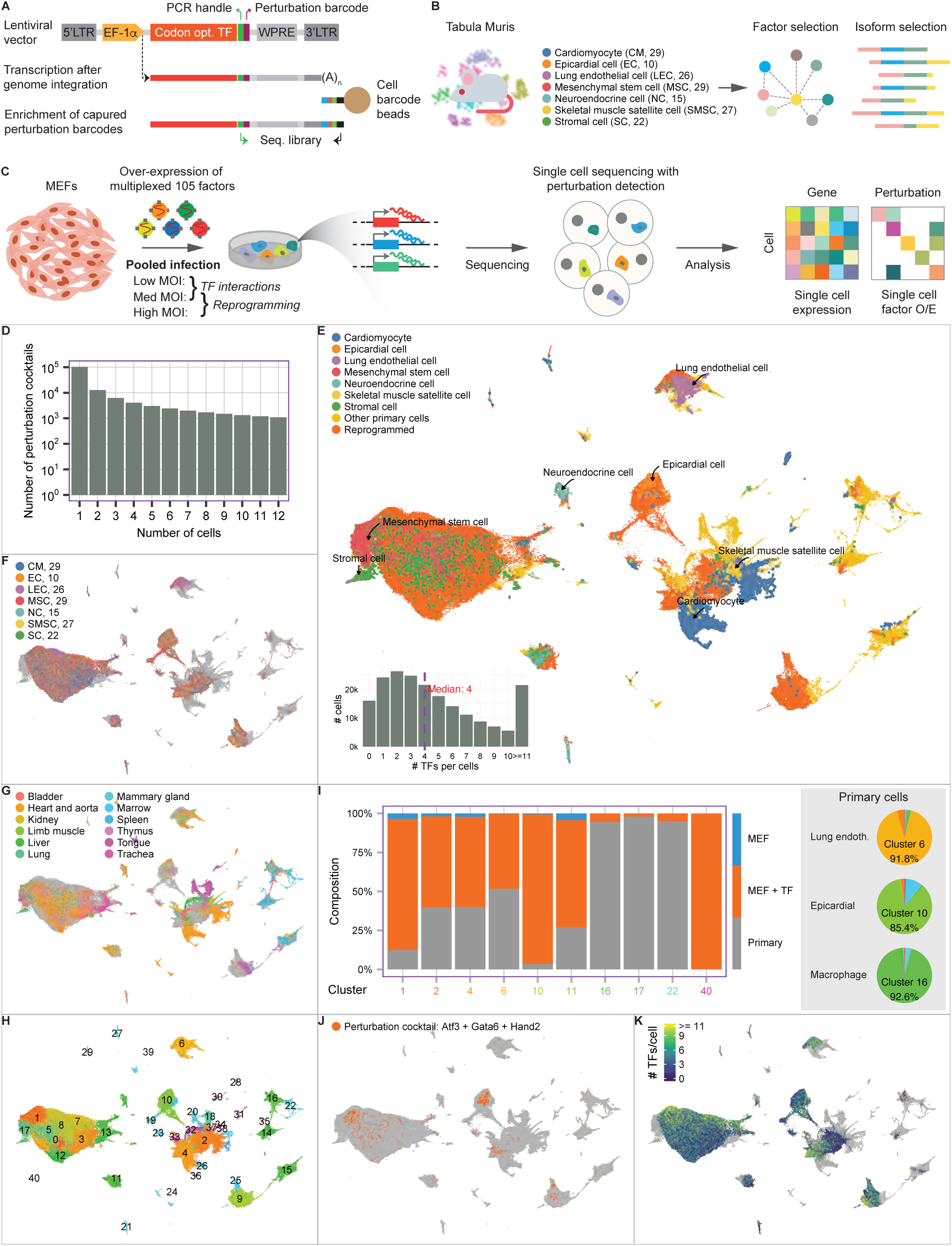
Diverse transcriptional states from combinatorial TF reprogramming. A. Schematic of Reprogram-Seq lentiviral TF ORF barcoding system compatible with single-cell RNA-Seq detection. B. Schematic of Reprogram-Seq applied to Tabula Muris cell types. For 7 cell types, 10 to 29 candidate TFs were prioritized for combinatorial over-expression studies. C. For each of the 7 cell type batches, MEFs were infected with lentiviral libraries for the candidate TFs, followed by single-cell RNA sequencing and analysis. D. Distribution of the number of cells recovered for distinct TF cocktails. E. UMAP visualization of reprogrammed MEFs and reference cell types. Reference cells are colored by cell type. Reference cell types (excluding “Other primary cells”) are plotted as larger dots to improve visibility. (insert) Distribution of the number of TFs identified in sequenced cells. F. UMAP feature plot, with cells colored by reprogramming batch. G. UMAP feature plot, with reference cells colored by origin tissue. MEF-derived cells are grey. H. UMAP feature plot, with cells numbered and colored by cluster. I. (left) For each cell cluster indicated, shown is the origin of cells in the cluster (control MEF, reprogrammed MEF, primary cell). The “Primary” label on the bottom indicates the most frequent primary cell type in the cluster. (right) For each primary cell type indicated, the pie chart shows the distribution of the primary cells across the clusters. J. UMAP feature plot, where cells receiving the epicardial TF combination Atf3 + Gata6 + Hand2 are colored orange. K. UMAP feature plot, with cells colored by the number of distinct exogenous TFs detected.

To apply Reprogram-Seq in a cost-effective way, we sought to minimize the number of TFs synthesized while maximizing the cell types where these TFs had reprogramming potential. Using Tabula Muris as a reference, we identified 105 cell-type specifically expressed TFs spanning 7 diverse target cell types (**Figure 1B**): cardiomyocytes (CM, 29 TFs), epicardial cells (EC, 10 TFs), lung endothelial cells (LEC, 26 TFs), mesenchymal stem cells (MSC, 29 TFs), neuroendocrine cells (NC, 15 TFs), skeletal muscle satellite cells (SMSC, 27 TFs), and stromal cells (SC, 22 TFs) (**Supplemental Table 1**). This synthesized set of TFs represents ∼7% of TFs encoded in the mouse genome.

For each of the 7 target cell types, we applied Reprogram-Seq to combinatorially overexpress a subset of the 105 TFs relevant to its transcriptional state (**Supplemental Table 1**) in mouse embryonic fibroblasts (**MEFs**). We refer to each of these 7 experiments as a batch. We infected MEFs across a range of lentiviral titers (low titer averaging 1-2 TFs per cell; high titer averaging 5-6 TFs per cell), to enable different downstream applications (**Figure 1C-E**). After 14 days of lentiviral infection, we performed single-cell RNA-Seq (scRNA-Seq) to detect the transcriptomes and TF perturbations in individual cells. Overall, we recovered ∼200k high-quality MEF-derived cells, with a median of 4 TFs per cell (**Supplemental Table 2**, **Figure S1C**). This dataset spans >100k unique TF combinations, with 1335 TF combinations having at least 10 cells (**Figure 1D**). As a negative control, we also sequenced MEFs infected with lentiviruses expressing fluorescent proteins instead of TFs (**Figure S1D-E)**.

### Combinatorial TF over-expression reprograms MEFs to diverse states

To examine how combinatorial over-expression of TFs alters transcriptional states, we compared MEF-derived cells to reference mouse atlases ^16,43^ (**Figure 1E**). By co-embedding the cells, we observed that reprogrammed MEFs (**Figure 1F**) transcriptionally cluster with diverse *in vivo* cell states (**Figure 1G**). 13 out of 41 clusters contained significant enrichment of perturbed cells, with an average of 4-fold enrichment over unperturbed MEFs (**Figure 1H-I**). These clusters contained 41% of all perturbed MEFs. To highlight the single-cell resolution of the TF perturbations, we examined known reprogramming cocktails. For example, we previously identified Atf3, Hand2, and Gata6 (3F) as an epicardial-reprogramming cocktail. Using Reprogram-Seq, we find that cells perturbed with 3F are significantly enriched in the epicardial cluster (p < 2.2e-16) (**Figure 1J**). Notably, consistent with the observation that known reprogramming cocktails often contain multiple TFs, the number of TFs induced in MEFs correlates with the frequency of transcriptional reprogramming (**Figure 1K**). Overall, these results suggest that our strategy effectively identifies TF combinations that perturb global transcriptional states.

We next analyzed the reprogramming outcome of each batch. First, for each cluster that represents a primary cell type, we examined the batch specificity of the reprogrammed cells in the cluster. That is, are all reprogrammed cells derived from the expected batch (e.g., reprogrammed CMs from the CM batch) or do some of them come from ‘non-specific’ batches (e.g., reprogrammed CMs from non-CM batches)? On one extreme, Cluster 6 represents cells derived from the LEC Reprogram-Seq experiment (which contained 26 TFs targeting lung endothelial cells) and 91.8% of all primary cells endothelial cells are in this cluster, demonstrating high batch specificity (**Figure 1I**). Unexpectedly, other clusters had almost even representation of cells from multiple batches of Reprogram-Seq experiments, even though each batch contains different TFs (**Figure 1G**). Second, we considered the specificity of the reprogramming outcome of each batch. Consistent with the results above, we find that one reprogramming batch often yields cells in multiple transcriptional states (**Figure S1G**) (19 transcriptional states per batch on average). For example, while MEFs induced with CM TFs were significantly enriched in the cardiomyocyte cluster, they were also enriched in clusters representing skeletal muscle satellite cells (**Figure S1F-G)**. Third, we also surprisingly find that perturbed MEFs are significantly enriched for unexpected cell states that were not intended targets for transcriptional reprogramming (**Figure 1I**). Notable examples include Clusters 16 and 22, which contain distinct hematopoietic cell types including macrophages (Cluster 16, p = 8.15E-102) and granulocytes (Cluster 22, p = 4.69E-42). These clusters are 5.5-fold more enriched in perturbed cells over control MEFs.

Overall, these results indicate that combinatorial TF induction reprograms MEFs towards diverse transcriptional states, many of which have similarity to in vivo cell states. Our observations highlight that diverse TF combinations can reprogram MEFs to similar transcriptional states. For instance, 16 out of 26 predicted TFs for lung endothelial cell reprogramming are enriched in cluster 6. These results also demonstrate that some TF combinations can reprogram towards unintended cell fates, which is consistent with documented examples of TFs with promiscuous activation of cellular regulatory programs ^44–47^.

### TF combinations drive cell-type specific regulatory programs

While clustering analysis can identify global transcriptional changes, it does not distinguish the gene regulatory programs driven by different TF combinations. To address this, we first identified 1335 distinct TF combinations (**Figure 2A**, **Supplemental Table 3**) that met stringent quality filters. This set consists of 104 TF singles, 801 TF doubles, 322 TF triples, and 107 combinations with 4 or more TFs (**Figure 2B**). We performed MDE embedding with parameters tuned to distinguish subtle transcriptional changes (**Figure 2C**) ^48^. In this embedding, different TF combinations occupied distinct transcriptional spaces. Many TF combinations were represented by multiple reprogramming batches (e.g., TF Lmo2 in CM, LEC, NC, SC, EC batches), suggesting that batch effects are limited (**Figure 2D-E**). We then used the density-based HDBScan method to identify 51 perturbation clusters, each consisting of multiple TF combinations (**Figure 2F**, **Figure S2A**). Clusters also contained distinct TF combinations from the same reprogramming batch (**Figure 2D**). For example, in Perturbation Cluster 6 (**Figure S2A)**, 81.8% of TF combinations are derived from the MSC reprogramming batch. In contrast, Perturbation Cluster 1 represents experiments from almost all reprogramming batches.

**Figure 2:**
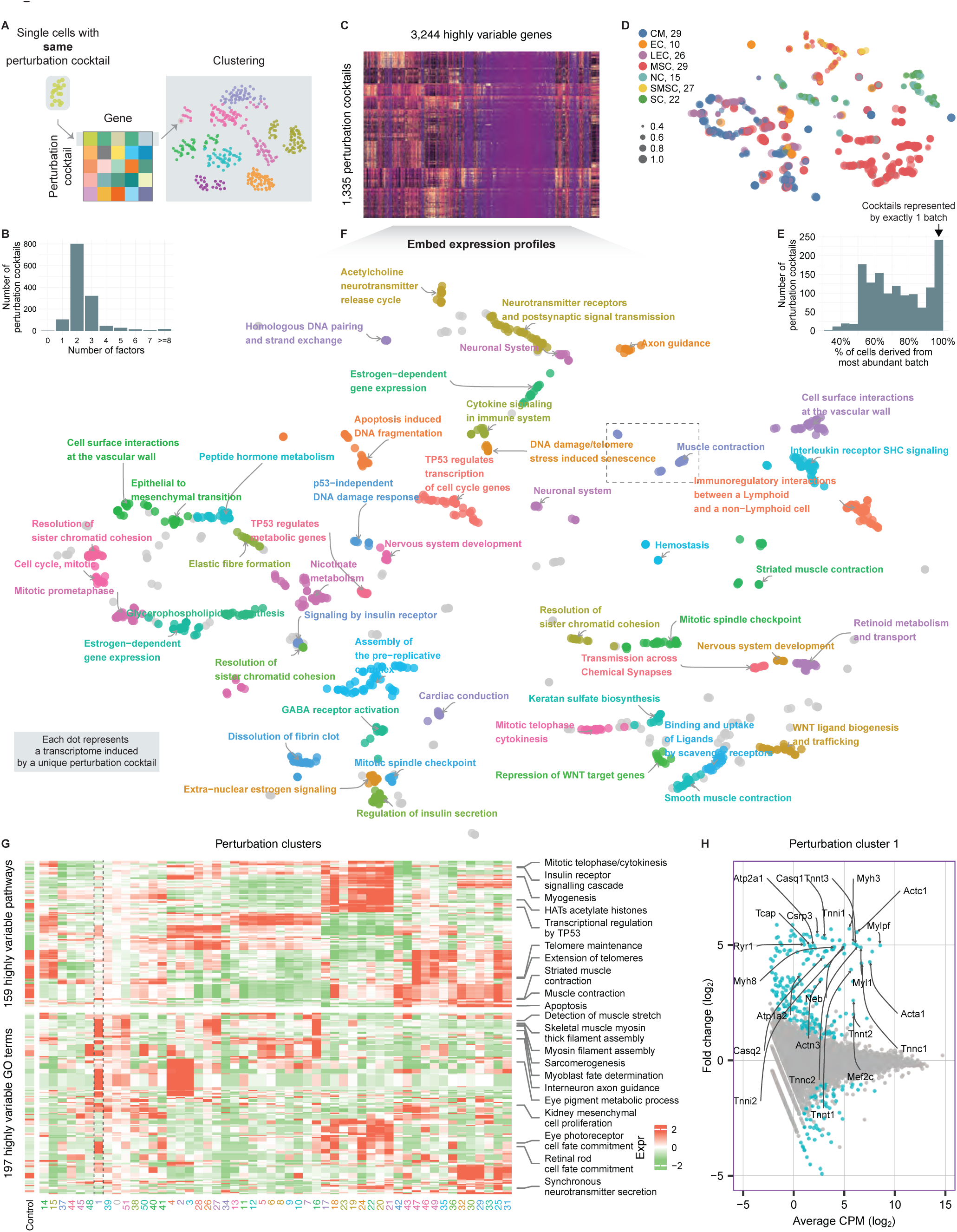
TF combinations drive cell-type specific regulatory programs. A. Schematic to visualize transcriptional phenotypes from combinatorial TF over-expression. Cells induced with identical TF combinations were combined. B. For the 1335 TF combinations analyzed, shown is the distribution of the number of TFs in each combination. C. The heatmap indicates the expression of 3244 variably expressed genes across the 1335 TF combinations. D. MDE embedding of transcriptomes for 1335 TF combinations. Each dot indicates a TF combination, colored by experimental batch. When a given TF combination is recovered by multiple batches, the size of each dot indicates the fraction of cells derived from the more abundant batch. E. Across the 1335 TF combinations, shown is the fraction of cells derived from the most abundant batch. F. MDE embedding, with TF combinations colored by perturbation cluster and annotated with enriched Gene Ontology terms. G. Heatmap illustrating the enrichment of Gene Ontology terms and pathways across the 51 perturbation clusters. H. The MA plot indicates the differentially expressed genes in perturbation cluster 1 (teal). Several genes with cardiac roles are indicated.

Perturbation clusters represent TF combinations that share similar transcriptional impacts (**Fig. 2F**). Since TFs drive gene regulatory networks (**GRNs**), the set of genes induced and repressed by a given combination represents an experimentally estimated GRN. To establish these GRNs, we identified the genes induced and repressed in perturbation clusters (**Supplemental Table 4**). To evaluate the biological relevance of these GRNs, we annotated them with several functional databases including Reactome, Gene Ontology, and KEGG (**Figure 2F-G**, **Supplemental Table 5**). One notable example is Perturbation Cluster 1, which induced genes highly enriched for cardiac function, including muscle contraction (Reactome ID: R-MMU-397014, p = 2.01E-28) and skeletal muscle myosin thick filament assembly (GO ID: GO:0030241, p = 3.8E-05) (**Figure 2G**). Consistently, well-known cardiac genes were highly induced in Cluster 1. For example, we observed a 3.8-fold induction of Tnnt2 (p = 1.13E-06) and a 15.7-fold induction of Tnnt3 (p = 3.87E-12) (**Figure 2H**). Thus, TFs in Perturbation Cluster 1 induce a GRN representing a muscle gene expression program. We also identify other candidate cell-type specific GRNs. These include GRNs that have enrichment for: nervous system development (Cluster 6, p = 2.04E-03), axon guidance (Cluster 18, p = 4.48E-04), Estrogen−dependent gene expression (Cluster 22, p = 2.52E-03), striated muscle contraction (Cluster 16, p = 1.17E-22), signaling by insulin receptor (Cluster 37, p = 5.32E-03), and cardiac conduction (Cluster 34, p = 2.20E-02). These analyses demonstrate that TF combinations can drive diverse cell-type specific gene programs (**Figure 2F**, **Figure S2B)**.

### TF combinations drive modular gene regulatory networks

The previous analysis focused on TF-induced GRNs at the level of perturbation clusters (containing multiple TF combinations) and pathways (containing multiple genes). To gain more granular knowledge of the GRNs, we next performed analysis at the level of distinct TF combinations and genes (**Figure 3A**).

**Figure 3:**
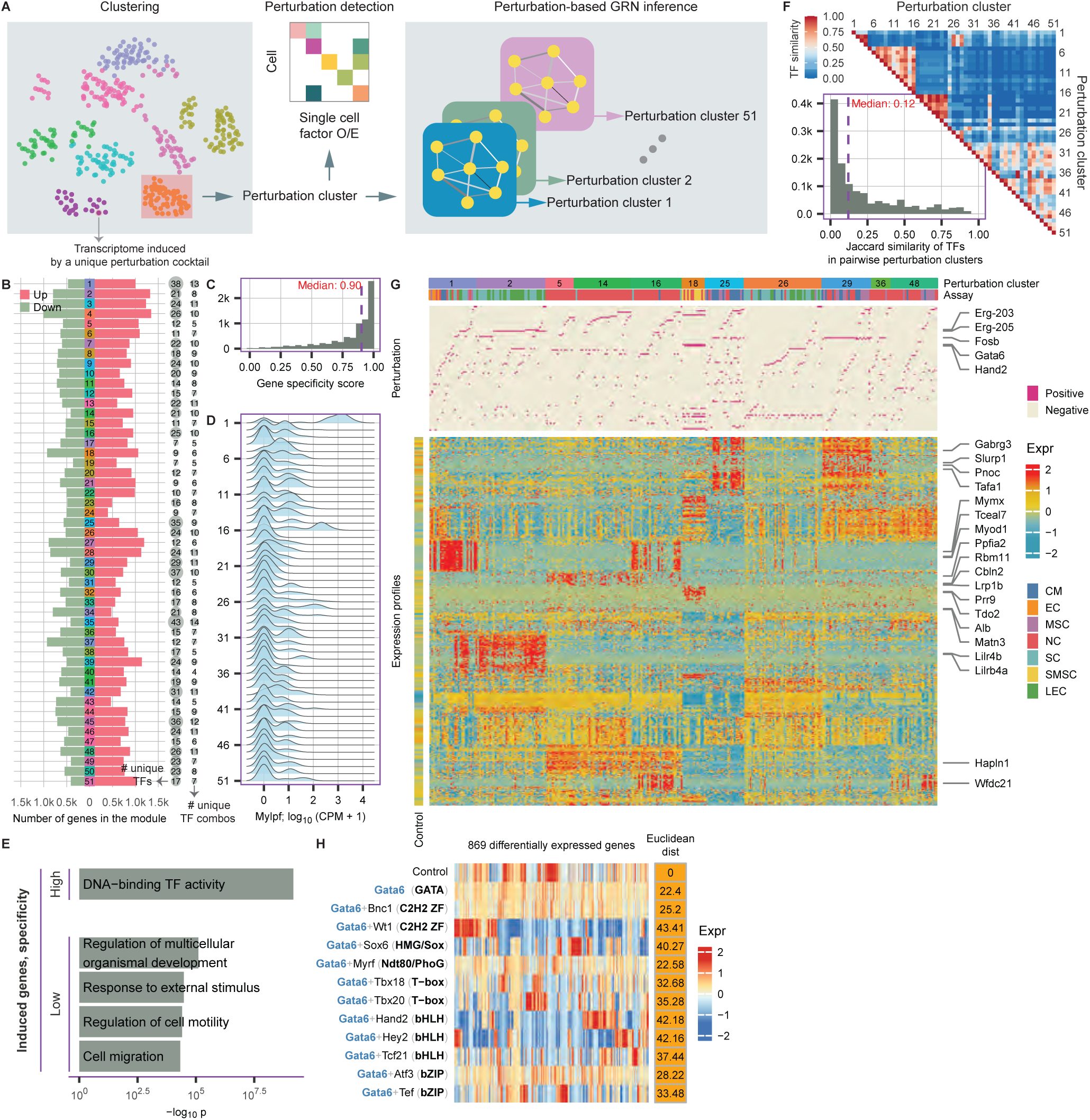
TF combinations drive modular gene regulatory networks. A. Schematic to estimate gene regulatory networks driven by perturbation clusters. Cells in each perturbation cluster are grouped, followed by differential expression analysis. B. For each perturbation cluster, shown is the number of differentially expressed genes induced and repressed. (left) The number of unique TFs in the perturbation cluster. (right) The number of unique TF combinations. C. Across all differentially expressed genes identified in B, shown is the specificity across modules (1 - fraction of occurrence across modules). A specificity score of 0 indicates a gene is found in all modules. D. Expression of Mylpf in cells across the 51 perturbation clusters. E. Gene ontology enrichment for high specificity and low specificity genes. F. The heatmap indicates the overlap fraction of TFs across perturbation clusters. The histogram quantifies Jaccard similarity of TFs across perturbation clusters. G. Heatmap indicating gene expression patterns (bottom) across perturbation clusters. Each column indicates a distinct TF combination (middle), grouped by batch/assay and perturbation cluster (top). H. The heatmap illustrates the expression of 869 differentially expressed genes in cells induced with Gata6 alone or Gata6 + TF_X_ compared to control MEFs. TF_X_ spans 11 different TFs from different TF families (bold). For each Gata6 + TF_X_ pair, the values on the right column indicate Euclidean distance relative to control cells.

First, to gain insights into individual genes, we examined the set of commonly induced and repressed genes in each perturbation cluster. On average, each perturbation cluster significantly induced 834 genes and repressed 607 genes (**Figure 3B**). The union across all perturbation clusters spans a total of 7751 differentially expressed genes. Surprisingly, despite the sparse sampling of the complete perturbation space (∼7% of all TFs, ∼7% of all single TF perturbations, and ∼0.04% of all double TF perturbations), we find that the TF-induced GRNs span ∼23% of all genes in the transcriptome. This observation suggests that sparse subsampling of combinatorial TF perturbations could be used to span transcriptome-wide gene regulation.

Next, to examine specificity, we assessed how often a given gene was recovered across GRNs. We observed that TF-induced GRNs highly overlap: 77.6% of genes were induced in multiple perturbation clusters (**Figure 3C**, **Figure S3A**). For example, Mylpf was highly induced in both perturbation clusters 1 and 16 (**Figure 3D**). Overall, each induced gene was found in a median of ∼5 perturbation clusters (0.90 specificity score; where 0 indicates a gene found in all modules). Genes with the lowest specificity were significantly enriched for Gene Ontology terms including Regulation of multicellular organismal development (p = 7.9e-06) and Response to external stimulus (p = 3.3e-05) (**Figure 3E**), indicating that the common features of GRN rewiring are similar to developmental responses. In contrast, genes with the highest specificity and induced in a small number of perturbation clusters are significantly enriched in DNA-binding TF activity (p = 6.7e-10), suggesting the activation of cell-type specific genes. Overall, these observations imply a shared regulatory structure of GRNs, and that multiple TF perturbations can be used to induce a given gene.

To examine the diversity of TF perturbations, we compared the composition of TFs across perturbation clusters (**Figure 3F**, **Figure S3B**). Supporting our observation that the transcriptomes induced by different clusters are distinct, we found that TF perturbations were also generally distinct across perturbation clusters. Notably, 25.0% of all perturbation cluster pairs share no TFs, and most pairs (61.6%) have very low TF overlap (Jaccard similarity < 0.2) (**Figure 3F**). These results suggest that perturbation clusters span diverse sets of TFs.

The observations above indicate a striking modularity of TF-driven GRNs at the level of individual TF cocktails and genes. **Figure 3G** illustrates an example (**Figure S3C**). Perturbation cluster 1 induces a specific module of cardiac and muscle genes. These genes are rarely induced by other perturbation clusters. Perturbation cluster 1 has 27 distinct TF cocktails spanning 38 unique TFs evaluated in 6 batches of Reprogram-Seq experiments. In contrast, Perturbation cluster 2 induces a module of endothelial marker genes, including Lilr4b and Lilrb4a, that is distinct from Perturbation cluster 1. Notably, the TF combinations in Perturbation cluster 2 are highly enriched for the ETS transcription factor Erg (Erg-205, 61.5%; Erg-203, 20.5%) or Fli1 (25.6%), suggesting that these TFs induce an endothelial gene program. Interestingly, a small set of TF combinations in perturbation clusters 1 induce both cardiac and endothelial gene programs. Highlighting the modularity of these gene expression programs, these TF combinations are also enriched for Erg and Fli1. Consistent with these observations, the ETS family of TFs are well-established lineage-specifiers of endothelial cell fate, and lineage tracing studies of Isl1+ cardiac progenitors demonstrate a common origin of cardiomyocytes and endothelial cells ^49,50^, and Nkx2-5 transactivates Etv2 in multi-potent progenitors cells to regulate endothelial/endocardial cell fate ^51^.

The analyses above indicate that TF combinations exhibit highly modular activities. To illustrate an example, we focused on the epicardial TF Gata6 and its interactions with 11 other TFs. Several TFs (Bnc1, Myrf, Atf3), when paired with Gata6, exhibit similar effects on the transcriptome compared to Gata6 induction alone (**Figure 3H**, **Figure S3D**). In contrast, pairing the remaining 8 TFs with Gata6 yields transcriptional responses that are distinct from Gata6 alone. Notably, each of these pairs Gata6 + TF_X_ induces a unique module of genes. For example, Gata6 + Wt1 induces an expression program most similar to primary epicardial cells, including the epicardial genes Bnc1, Gpm6a, Myrf and Upk1b. Interestingly, we observe distinct behaviors of Gata6 + TF_X_ pairs even when TF_X_ belong to the same family of transcription factors, suggesting diverse functions for TFs that share structural similarities at the level of protein domains, and often binding motifs (**Figure S3E**).

### Top-down analysis nominates TF cocktails that reprogram transcriptional state

One goal of Reprogram-Seq analysis is to identify TF cocktails that reprogram transcriptional state. In a previous study, we used pseudotime analysis to nominate a cocktail of 3 transcription factors (Atf3, Gata6, Hand2; AGH) for epicardial reprogramming. Since the current study has greater resolution of cellular perturbations, we first revisited epicardial reprogramming with the goal of systematically ranking TF combinations. Pseudotime analysis illustrates a cellular trajectory from control MEFs to primary epicardial (Krt19+) cells that is spanned by reprogrammed MEFs (**Figure 4A**, **Figure S4**). Across the pseudotime trajectory, reprogrammed MEFs induce different signature genes, culminating in the endogenous activation of epicardial genes including Gpm6a, Wt1, and Krt19 (**Figure 4B-C**).

**Figure 4:**
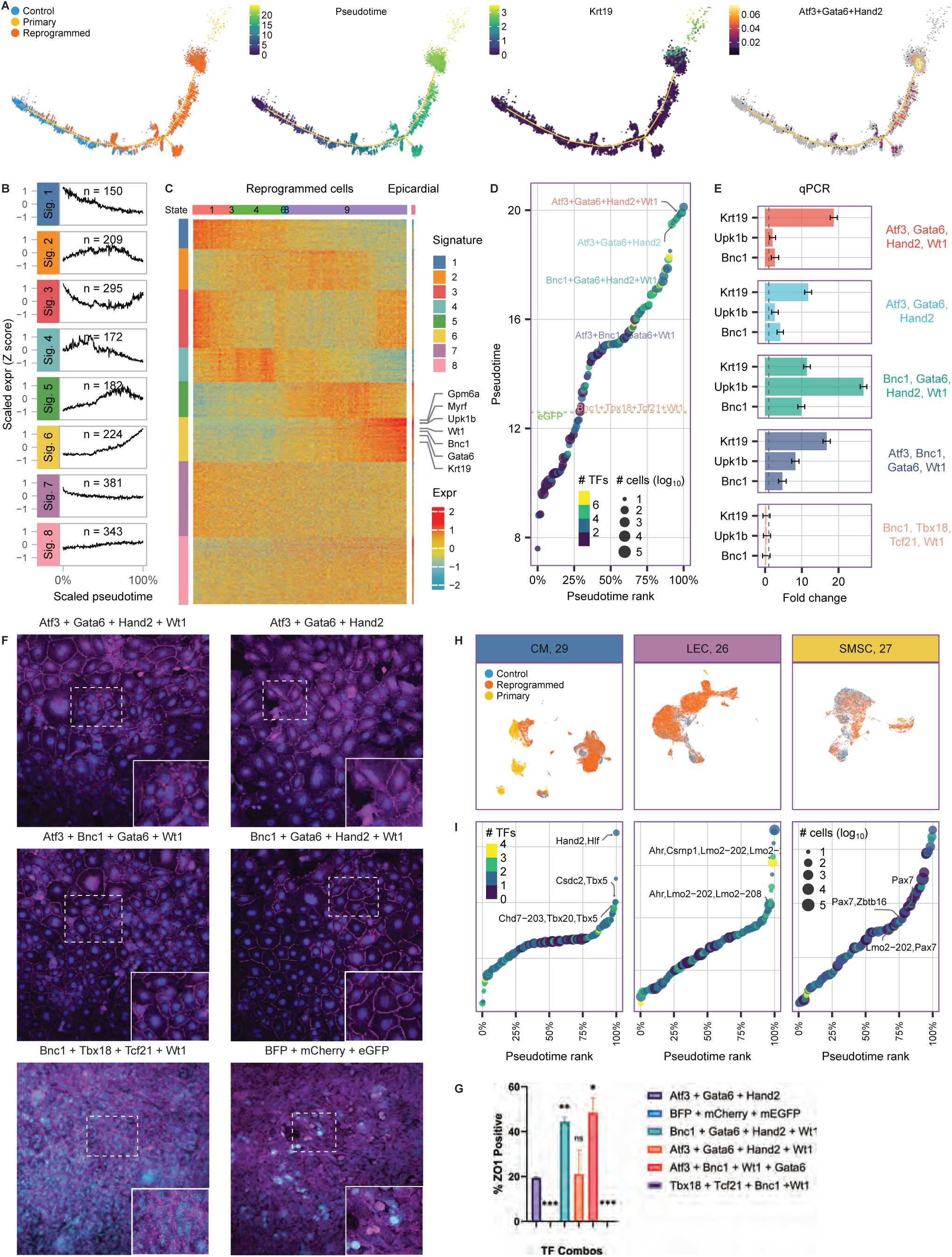
Prediction of TF combinations for transcriptional reprogramming. A. Pseudotime trajectory of epicardial reprogramming. (left) Indicated are control MEFs (blue), TF-induced MEFs (orange), and primary epicardial cells (yellow). (middle left) Feature plot colored by pseudotime. (middle right) Feature plot colored by the expression of epicardial marker gene Krt19. (right) Feature plot colored by cells induced with Atf3 + Gata6 + Hand2. B. Clusters of distinct gene expression patterns across epicardial transcriptional reprogramming. Pseudotime is indicated by the x-axis. C. The heatmap shows the expression of the gene clusters in C across all MEFs induced with epicardial TFs (ordered by pseudotime). D. Distinct TF combinations are ordered by transcriptional reprogramming performance according to pseudotime. Several TF combinations are indicated. Atf3 + Gata6 + Hand2 was previously identified as an epicardial reprogramming cocktail. E. Bulk qPCR validation of putative TFs for transcriptional reprogramming of epicardial cells. F. Immunostaining of tight junction protein ZO1 (a feature of epicardial cells), in MEFs after over-expression of putative TFs for epicardial reprogramming. G. Quantification of ZO1 positive cells in panel F. H. Across all reprogramming batches and targeted primary cells, shown is the clustering of control MEFs (blue), MEF-derived cells (orange), and primary cells (yellow). I. Pseudotime analysis to rank putative TFs for neuroendocrine and stromal cell reprogramming.

Consistent with our observations above (**Figure 1**), we find that cells receiving more TFs generally exhibited better performance in transcriptional reprogramming compared to cells with fewer TFs (**Figure 4D**, **Figure S4B-C**). However, this trend does not appear to be additive, as cells with the highest number TFs (>6) only exhibited moderate performance. These observations indicate the utility of cells with moderate to high titer in identifying novel reprogramming cocktails. In agreement with our previous study, we also observed that cells co-induced with AGH are highly enriched towards the end of pseudotime (93.3th percentile). Notably, we identify several TF combinations with pseudotime performance that exceeds that of AGH, including a 4-TF combination AGHW (AGH + WT1) (97.5th percentile). Consistent with this role in reprogramming, WT1 (Wilms’ tumor 1) is a highly expressed transcription factor in developing epicardial cells and is an essential regulator of epicardial cell development ^52,53^. Independent qPCR experiments validated newly identified TF cocktails for transcriptional reprogramming towards epicardial states (**Figure 4E**). Finally, to investigate reprogramming phenotypes outside of transcription, we examined the localization of the tight junction protein ZO1, which is a feature of epicardial cells. Confirming our approach, we observe that MEFs induced with the newly identified TF combinations exhibit strong ZO1 localization at cell-cell junctions (**Figure 4F-G**).

To test the generality of this approach, we extended this analysis to the remaining batches of Reprogram-Seq experiments (**Figure 4H-I, Figure S4D-E**). By anchoring primary cells as a destination transcriptional state, we ranked TF cocktails for transcriptional reprogramming. For example, in the LEC experiment, this analysis recovers Aryl hydrocarbon receptor (Ahr) as a strong TF in several high-ranking TF cocktails. Ahr has an essential role in maintaining the function of lung endothelial cells during viral infection ^54^. In the smooth muscle stem cell (SMSC) experiment, one TF appearing in several high-ranking combinations is Pax7, which has been documented as an important reprogramming factor for satellite cells ^55^. Finally, while we do not recover cells with simultaneous expression of the well-known cardiac reprogramming factors such as Gata4, Hand2, Mef2c, and Tbx5 in the CM experiment (because the combinatorial space was too large), we did recover enrichment of individual factors in high-scoring cocktails ^56^.

### TF-TF interactions influence reprogramming potential

The basic unit of combinatorial activity is a pair of TFs. Several modes of TF interaction include independence, dominance, and cooperativity ^37,57–59^. To mathematically define how TF pairs functionally interact in the context of transcriptional reprogramming, we used a previously established framework to model interactions by linear regression ^37^. Briefly, we modeled changes in gene expression observed in doubly perturbed cells (AB) as a linear combination of singly perturbed cells (A and B alone) (**Figure 5A**):

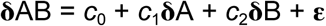

**Figure 5:**
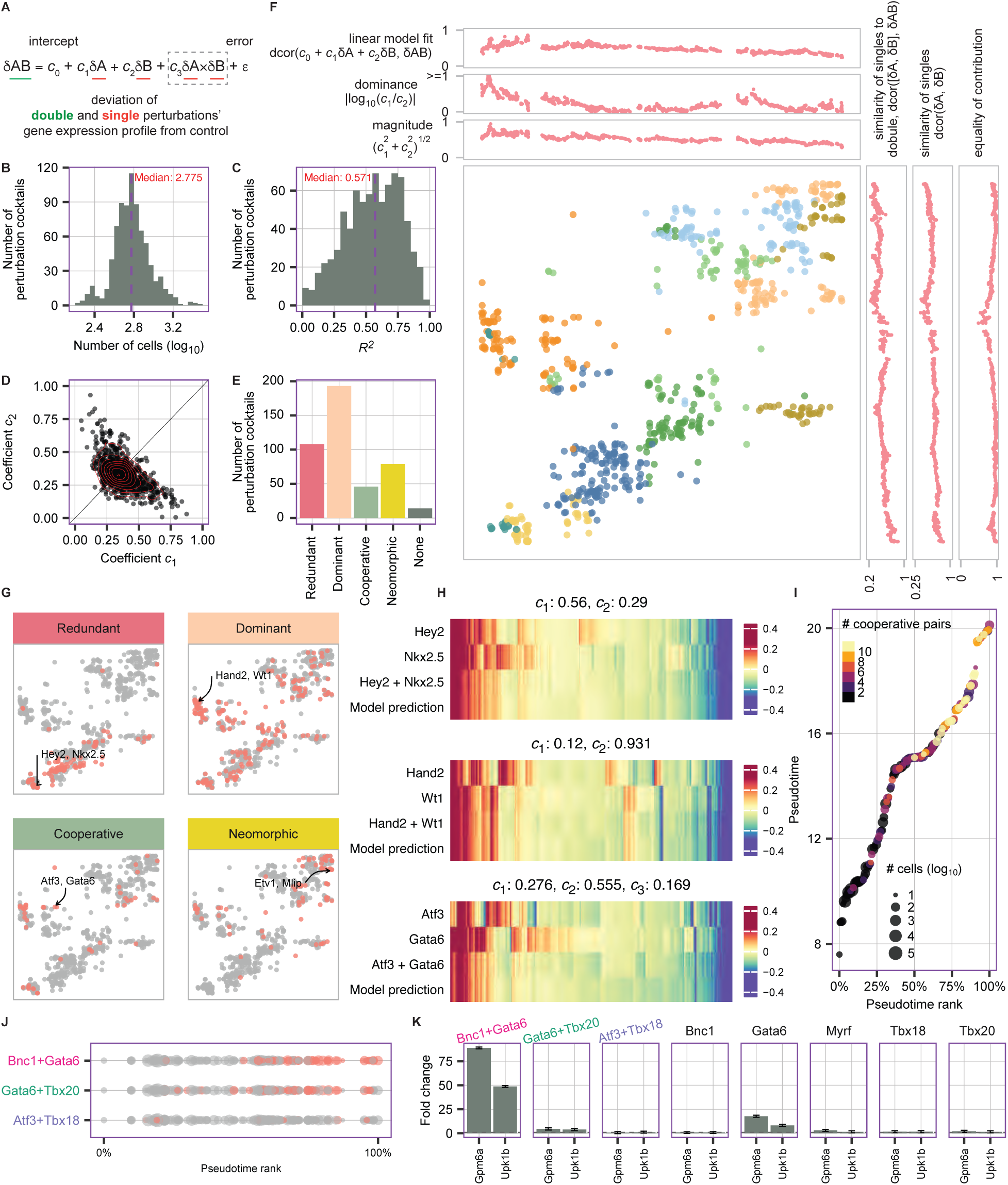
Transcription factor cooperativity in transcriptional reprogramming. A. A linear model to decompose combinatorial TF perturbations from single TF data. The c_1_*δa* and c_2_*δb* terms model independent contributions of individual TFs A and B, while the c^3^ x *δa* x *δa* term models cooperativity. B. Distribution of *R^2^* across linear models across 801 TF pairs. C. Across all 801 TF pairs, shown is the distribution of *c_1_* and *c_2_* values in the linear model. D. Distribution of the number of cells used in each application of the linear model. E. Visual embedding of TF pairs based on parameters of the linear model. F. The bar chart indicates the number of TF pairs with the following relationship between constituent TFs: redundant, dominant, cooperative, neomorphic, and none. G. Feature plots indicating TF pairs with redundant, dominant, cooperative, and neomorphic relationships between constituent TFs. Example TF pairs are indicated. H. Example TF pairs with redundant (Hey2 + Nkx2.5), dominant (Hand2 + Wt1), and cooperative (Atf3 + Gata6) relationships. For each of the three plots, shown is the transcriptome expression pattern for: cells induced with each individual TF (top 2 rows), cells induced with both TFs (third row), and the prediction from the linear model (bottom row). The model parameters are indicated above each plot. I. TF interaction relationship in the context of epicardial reprogramming. J. For the epicardial reprogramming batch, cells were ranked by pseudotime and colored by whether they contain the indicated TF pairs. K. qPCR analysis of Gpm6a and Upk1b expression for pairwise and individual TFs.

Here, c_1_**δ**A and c_2_**δ**B represent linear effects and **ε** is an error term that encapsulates non-linear effects. We applied this linear model to 801 TF pairs. 14 pairs were excluded from the downstream analysis (F-test, p >= 0.05, suggests this model doesn’t work). On average, 595 cells were used for modeling (median, 10 ^ 2.775) (**Figure 5B**). Overall, the linear model performed well, with most tested TF pairs yielding R^2^ values above 0.5 (median 0.571) (**Figure 5C**, **Figure S5A**). This good linear fit was also supported by the distance correlation metric (dcor, **Figure S5B**) ^37^. Visual embedding of these linear model fit parameters indicate that TF interactions are diverse (**Figure 5F-G**, **Figure S5A-I**). To gain more insights into TF interactions, we developed statistical tests to identify several classes of interactions (see Methods). Notably, the most frequent mode of TF-TF interaction is dominance (24.5%) (**Figure 5E**, **Figure S5C-F**), where one TF masks the impact of another TF (e.g., Hand2 and Wt1; **Figure 5G-H**). Redundant interactions are the next most frequent (13.7%), where two TFs have similar transcriptional impacts, alone or in combination (e.g., Hey2 and Nkx2.5; **Figure 5G-H**, **Figure S5G**).

Published studies indicate that reprogramming TFs can interact nonlinearly ^25,37^. To represent one type of non-linear cooperative interaction between A and B, we augmented our linear model by incorporating a cooperative component (*c*_3_**δ**A⨯B), similar to a previous study ^17^ (**Figure 5A**, **Supplemental Table 6**):

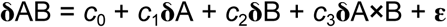

Using this augmented linear model, we identified 46 TFs that behaved cooperatively (*c*_3_, p < 0.05), where TFs are multiplicatively cooperative (e.g., Atf3, Gata6; **Figure 5G-H**). To characterize other types of non-linear interactions, we also identified 79 neomorphic interactions (large mean absolute error). (e.g., Etv1 and Mlip; **Figure 5G, Figure S5H**).

TF-mediated reprogramming is the outcome of multiple TFs interacting to reconfigure the transcriptome. We wondered if linear (redundant, dominant) or non-linear (cooperative, neomorphic) interactions were more likely to lead to transcriptional reprogramming. To address this question, we examined the enrichment of linear and non-linear interactions in TF cocktails as a function of transcriptional reprogramming (as defined by pseudotime). We observed that TF-TF interactions from successfully reprogrammed cells are significantly enriched in cooperative interactions (one-tailed *t*-test, p-value = 3.44e-14) (**Figure 5I**, **Figure S5J)**. One example of a cooperative TF cocktail for epicardial reprogramming is Bnc1 and Gata6 (*c*_3_ = 0.14, p-value = 7.05E-08). To test if this TF pair exhibits non-linear behavior, we compared with two other TF pairs from the epicardial batch (Gata6 + Tbx20; Atf3 + Tbx18; **Figure 5J**). qPCR experiments confirm that, while Bnc1 or Gata6 alone do not induce the epicardial genes Gpm6a and Upk1b, the combination of both TFs induces non-linear induction of these target genes (**Figure 5K**).

### Predicting TFs driving single-cell transcriptional phenotypes

The analysis in Figure 4 represents a top-down approach using cells that have reprogrammed to a transcriptional state resembling *in vivo* cells to gain insights into reprogramming TFs. However, this approach requires reprogrammed transcriptional states to be well-represented by perturbed MEF-derived cells. As an alternative approach, we next developed a bottom-up approach to predict higher-order TF cocktails from simpler TF combinations.

To test the feasibility of this approach, we first focus on the simpler problem of predicting the identity of single TF perturbations from single-cell transcriptomes (**Figure 6A**). We used two approaches, a linear model with stochastic gradient descent (SGD) learning and the XGBoost algorithm, to train the models using a filtered set of 66 single-TF perturbations well-represented by high quality sequenced cells (>= 100). To assess the accuracy of the models, we used a computational cross-validation approach, with 80% of the dataset used for training and 20% withheld for testing. Since the results of the two approaches are comparable (**Figure S6A**), we only refer to the results of the linear model in the main text. Overall, we observe that single-cell transcriptomes have enough information to predict individual TF perturbations (**Figure 6B**). On average, 24.5% of cells can be correctly classified into the 66 TF categories, compared to an expectation of 1.5% by random guessing (**Figure 6D**) (odds ratio 16.3). Supporting the robustness of this approach, we observed similar trends on a smaller set of 36 single-TF perturbations with more sequenced cells >= 200 cells, with an odds ratio of 14.8 (**Figure S6B**).

**Figure 6.**
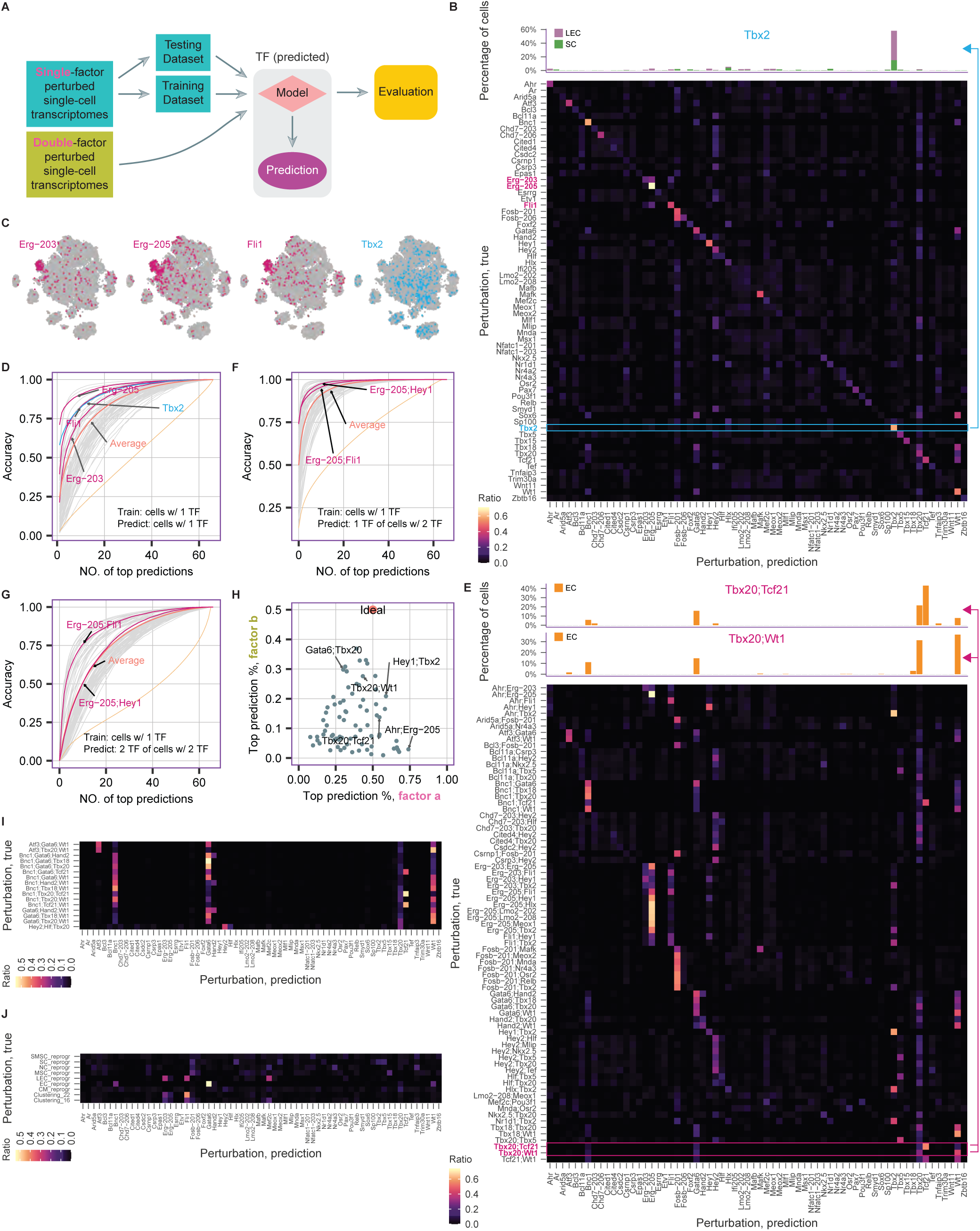
A. Schematic of approach to model transcriptional responses from single-TF experiments, and to use this model to predict TF perturbations from multi-TF experiments. B. (bottom) Heatmap illustrating the performance of a linear model to predict TF perturbations from transcriptional responses in cells with exactly 1 TF. Columns indicate predictions and rows indicate observations. Heatmap units are the fraction of cells in each row (which sums to 1). (top) TF predictions for cells with Tbx2. C. UMAP embedding of cells with exactly 1 TF. Feature plots indicate TF perturbations. D. Receiver operating characteristic (ROC)-like curves illustrating the performance of predicting single TF perturbations from single-cell transcriptomes, after training on cells perturbed with 1 TF. Dotted line indicates performance where TF labels are randomly shuffled. E. (bottom) Heatmap illustrating the performance of a linear model to predict TF perturbations from transcriptional responses in cells with exactly 2 TFs. (top) F. As in D, but illustrating the performance of predicting one of TFs of the double TF perturbations from single-cell transcriptomes, after training on cells perturbed with 1 TF. G. As in D, but illustrating the performance of predicting two of the TFs of double TF perturbations from single-cell transcriptomes, after training on cells perturbed with 1 TF. H. When predicting the perturbations of cells with 2 TFs (a and b), shown is the frequency that the top prediction is either a or b. I. As in (E), but predicting perturbations in cells that have exactly 3 TFs. J. As in (E), but predicting perturbations in cells that cluster near primary mouse cell types.

Several examples are illustrative. One class consists of predictions specific to one TF. For example 58.0% of cells induced with the T-box transcription factor Tbx2 are correctly predicted to be perturbed by the TF using each cell’s transcriptional profile alone (**Figure 6B**). A second class contains predictions specific to a small set of related TFs. For example, we tested two isoforms of Erg (denoted as Erg-203 and Erg-205), and the model occasionally predicts the incorrect TF isoform. For example, 25.4% of Erg-203 perturbed cells are predicted as Erg-203, while 3.66% of Erg-205 perturbed cells are predicted as Erg-203. Supporting the idea that these two isoforms likely induce similar transcriptional phenotypes, embedding of single-TF perturbed cells shows that Erg-203 and Erg-205 co-cluster while being distinct from other TFs like Tbx2 (**Figure 6C**, **Figure S6C** with all feature plots). Interestingly, Fli1-induced transcriptomes share similarity with those induced by Erg isoforms. Similarly, the misclassification of cells induced with Wt1 and Gata6 suggest similar transcriptional responses for these genes (**Figure 6B**, **Figure S6C**). In addition, we observe that while most cells with Nkx2-5 are correctly classified, a subset is also misclassified to Hey2. These observations could suggest similar transcriptional responses between these TFs. Consistent with this possibility, the TF interaction analysis indicates that Hey2 and Nkx2.5 have a redundant relationship (**Figure 5G-H**). This redundancy of transcriptional responses between distinct TFs complicates the computational task of correctly classifying cells by TF perturbations.

Next, we explored several parameters of the prediction model. At one extreme, we found that retraining the model on the top 5 performing TFs alone yielded 59% accuracy, compared with the bottom 5 performing TFs at 32% accuracy. We asked why some TFs were more difficult to predict than others. One possibility was that poorly predicted TFs have smaller gene expression changes. However, we observed no correlation between the number of differentially expressed genes induced by a TF and the ability to predict the identity of perturbed cells (**Figure S6D**). We also considered the possibility that the number of sequenced cells correlated with prediction performance. But we observed instances where highly sequenced TFs had poor performance and lowly sequenced TFs had high performance (**Figure S6E**, **Supplemental Table 7**). Motivated by the redundancy of TF effects observed above, we also considered the second-best or third-best predictions for the full set of 66 TFs (**Figure 6B,D**). For example, considering the top 3 predictions, performance increased to 41.3% accuracy, compared to 7.18% expected by chance. We next asked if removing redundant TFs improves prediction performance. To systematically test the impact of redundancy, we dropped each TF one at a time, retrained the model, and re-assessed performance. Globally, we only noticed a marginal improvement (24.7%) in prediction performance (**Supplemental Table 7**). However, some predictions significantly improved. For example, removing the transcription factor Fosb-201 yielded a 2.56-fold increase in prediction performance for Fosb-206. Overall, these results suggest that multiple factors will need to be considered to improve prediction performance in the future.

The analysis above predicted single TF perturbations from single-cell transcriptomes. We next asked whether this computational model could be applied to predict perturbations in cells receiving multiple TFs. We focused on 75 high quality TF pairs represented by > 50 cells (**Figure 6E-F**). Several examples are illustrative. First, some TFs (Bnc1, Erg-205, Fox-201, Hey2) are frequently found among the 75 TF pairs, and these are represented as vertical stripes of enrichment in the heatmap. Second, TFs with accurate prediction performance in the singlet TF analysis generally performed well in the doublet analysis. For example, Tbx20 and Tcf21 scored highly compared to Tbx18 in the singlet analysis. Likewise, prediction performance for doublets Tbx20-Wt1 and Tcf21-Wt1 was higher than Tbx18-Wt1. Third, some well-predicted singlet TFs are poorly predicted as doublets. For example, the singlet model poorly predicts Ahr when it is part of a TF doublet. Fourth, when other TFs like Fosb-201 are paired with another TF, only predictions for Fosb-201 are made. These last two examples may highlight dominance relationships between TF pairs (**Figure 6H**). Next, we asked whether both perturbations in a TF pair were represented in the top predictions (**Figure 6G**). On average, the model recovers 50.7% of TF pairs in the top 10 predictions, compared to 17.1% expected by chance.

Finally, we extended this analysis to cells with higher-order combinations of TFs. Although we only had sufficient cell numbers for 16 distinct TF triplets, the singlet model performed well to predict the TFs in these cells (**Figure 6I**). On average, we observed that top 3 TF predictions from the model matched the observed TFs 35.4% of the time. To further extend this analysis to predict TFs driving cellular reprogramming, we examined MEF-derived cells that co-clustered with in vivo cells (**Figure 1**). Consistent with our results above, the model identified Bnc1, Gata6, and Wt1 as TFs for reprogrammed epicardial cells (EC). It also highly scores Erg, Fli1, and Mef2C in endothelial cell reprogramming, consistent with the roles of these genes in maintaining endothelial cell fate and in angiogenesis ^60–62^. Finally, our reprogramming experiments in **Figure 1** also recovered cells that unexpectedly co-clustered with immune cells from Tabula Muris (clusters 16 and 22). Our computational model indicates that Erg, Fli1, and Mef2C are among the highest scoring TFs for these immune clusters. This result is consistent with the known roles of these TFs in maintaining homeostasis in hematopoietic lineages and blood cell reprogramming ^63–65^.

## DISCUSSION

One key finding of our study is that, despite a vast combinatorial space of TF combinations, the space of transcriptional outputs is much more limited. After overexpressing combinations of 105 TFs, clustering of single-cell transcriptomes shows that many cells induced with TFs cluster with *in vivo* cell types. Notably, the generation of *de novo* cell states was rare, at least on a transcriptome-wide level. At the finer resolution of gene modules, we identified ∼50 perturbation clusters. Surprisingly, while each cluster contains distinct TF combinations, they independently activate common gene modules. For example, Perturbation Cluster 1 contains diverse TF combinations that specifically induce a cardiac regulatory program (**Figure 3**). This striking modularity suggests a shared regulatory architecture of TFs on gene expression states. Together, these results support a many-to-few relationship between TFs and cell states. This simplifying organizational principle of TF-driven gene modules and gene regulatory networks has implications for cell state engineering and the topology of developmental regulatory programs.

TFs do not function in isolation. Traditionally, testing the cooperativity of a given pair of TFs required low throughput individual experiments ^66^, and recent technologies including single-cell screens have increased scale to moderate throughput ^25,37,67^. Motivated by a recent study using CRISPRi Perturb-Seq to knock down gene expression ^37^, our approach is to systematically define TF cooperativity through high throughput TF over-expression. Our observations highlight that, on a transcriptome-wide level, most TF pairs interact independently or redundantly. While comparatively fewer TF pairs have cooperative or neomorphic interactions, these interactions are over-represented in TFs that can successfully reprogram transcriptional state. These results suggest that complete sampling of all TF interactions may not be unnecessary and that focused sampling strategies could be used to map the interactions in an economical way.

GRNs comprise all regulatory inputs and transcriptional outputs for sets of functionally related genes ^9^. During embryo development, GRNs possess a strictly hierarchical structure. Upper level GRNs (“kernels”) establish the overall body plan through highly recursive and interconnected networks that do not tolerate change ^8^. Lower level GRNs, such as specification and differentiation GRNs, are much more adaptable and less recursively wired. During embryogenesis, sequential activation of specification and differentiation GRNs is coupled with canalization, the process by which alternative cell fates become restricted. Thus, direct reprogramming requires simultaneous de-canalization and alternative cell fate specification. In our framework, gene modules can be viewed as GRN sub-circuits that are re-deployed in multiple cellular contexts. Identification of which TFs activate a particular gene module may thus suggest how GRN topology is wired (or rewired). For example, the “muscle contraction” module is activated by many different TFs across several Reprogram-Seq batches. In contrast, the “blood vessel” module is dominantly activated by ETS family TFs. This striking difference exemplifies the unique GRN architectures that can exist. For the “muscle contraction” module, a horizontal or distributed topology likely reflects numerous cis-regulatory inputs capable of activation by a variety of TF combinations. For the “blood vessel” module, however, the vertical topology likely represents the dominant action of a single TF to specify cell fate, which may in turn reflect the distributed nature of endothelial cells within vertebrate organisms. In this way, analysis of Reprogram-Seq datasets enables reconstruction of GRNs using a top-down approach, which is distinct from traditional bottom-up dissection of regulatory element evolution.

The development of generalizable tools for the rational manipulation of cell type-specific transcriptional states will improve our understanding of the molecular basis of transcriptional states. Here, we focused on overexpressing the open reading frames of TFs, the most common approach for successful cellular reprogramming. The advantage of TF overexpression is the robustness of induction, which may be required to overcome the cell’s existing transcriptional state. Disadvantages include being exogenous, cellular integration by lentivirus, and non-physiological levels of induction. Similar approaches using CRISPRa have also recently been used for cellular reprogramming screens, which induce TFs endogenously. However, one disadvantage of this approach is that the level of TF induction is lower, and could be influenced by locus-specific epigenetic effects and/or chromatin accessibility. Alternatively, small molecule approaches have also been successfully applied to iPSC reprogramming, though they are currently less amenable to rational design and more difficult to customize gene targets ^68,69^. Another focus of our study is on the direct reprogramming of differentiated fibroblasts to other differentiated cell states. In contrast, parallel studies have focused on the “programming” of pluripotent stem cells to differentiated cell states, with similarities to differentiation during development. Both approaches have advantages and disadvantages. Advantages of direct reprogramming is the long history of studies using MEFs to derive diverse cell types and the ease of genetically manipulating MEFs. Advantages of programming human pluripotent stem cells is a more plastic epigenetic state and the ability to directly translate results for regenerative medicine. In sum, all of these resources will provide critical information needed to engineer transcriptomes for *in vitro* cell state modeling ^70–72^, with the aspirational goal of providing engineered cellular substrates for research and medicine ^73^.

## ACKNOWLEDGEMENTS

We acknowledge the BioHPC computational infrastructure at UT Southwestern for providing HPC and storage resources that have contributed to the research results reported within this paper. G.C.H is supported by CPRIT (RP190451), NIH (DP2GM128203, UM1HG011996, 1R35GM145235), the Burroughs Wellcome Fund (1019804), the Welch Foundation (I-2103-20220331), and the Green Center for Reproductive Biology. N.V.M. was supported by the NIH (HL136604, HL151650, UM1HG011996, and HG012768), the Burroughs Wellcome Fund (1009838), and the Department of Defense (PR172060).

## METHODS

### EXPERIMENTAL METHODS

#### Cell lines

Commercial Mouse Embryonic Fibroblasts (cMEFs) were purchased frozen from Cell Biologics, defrosted as needed, and grown in growth media (DMEM high-glucose + 10% FBS + 1% penicillin/streptomycin + 1% 100X GlutaMAX).

#### Construction of lentiviral vectors

TFs were codon optimized, synthesized, and cloned by Twist Bioscience into a lentiviral vector driven by EF1a and modified to contain 10X Genomics Capture Sequence 2 and a PCR handle (**Supplemental Table S8**) Each insert is composed of (5’ -> 3’): Kozak sequence (GCCACC), TF open reading frame (ORF), a common PCR handle, and an ORF-specific barcode. The eGFP-bc2 v1.3 plasmid map annotates elements in the vector (**Supplemental Table S8**).

#### Viral packaging, infection, and single cell reprogramming

A packaging cell line HEK 293Ts were used to produce lentivirus for reprogramming. Packaging cells were transfected using LT-1 per manufacturer’s instruction with lentiviral packaging vectors psPAX2 and pMD2.G in ratio of 2:1:3 with lentiviral vectors expressing specific transcription factors. Approximately 14-16 hours after transfection, the viral supernatant was aspirated and replaced with new media containing caffeine at a concentration of 2mM. 48 hours after transfection, virus was harvested, filtered (0.45um pore) and concentrated via ultracentrifugation (insert protocol reference here). The viral pellet was then resuspended in growth media (DMEM high-glucose + 10% FBS + 1% penicillin/streptomycin + 1% 100X GlutaMAX) plus 8ug/mL of polybrene (Sigma-Aldrich, St. Louis, MO) and applied to MEFs. The cells were incubated in growth media for 14 days and subsequently washed with DPBS and trypsinized into a single cell suspension. Cells were spun at 500 xg for 5 minutes at 4C and resuspended in cold growth media.

#### Viral titering

To determine the relative titer of each TF in cell type pools, bulk RNA-seq libraries amplifying only our exogenous perturbations were constructed. After extracting RNA using the Zymogen Quick RNA prep kit, Reverse Transcription was performed using an oligo containing the CS2 sequence from the 10X platform. After a PCR purification using the Qiagen Minielute kit, the cDNA is amplified using the 10X feature primer, then size selected using a 0.6X SPRIselect single sided selection. Another cDNA amplification on the PCR handle was performed, purified, and then amplified one last time to affix Nextera indices. A final 1.2X SPRIselect single sided selection was performed, then the libraries were tested for Quality Control metrics and sent off for next generation sequencing.

#### Flow cytometry

For flow cytometry, cells were trypsinized and resuspended in growth media. Cells were then spun at 500 xg for 5 minutes at 4°C and washed with room temperature DPBS. Cells were spun again at 500 xg for 5 minutes at 4°C and resuspended into a single-cell suspension in DPBS with 0.04% BSA. Flow cytometry was carried out on a Beckman Coulter CytoFLEX flow cytometer.

#### Cell hashing

We adopted the Cell Hashing protocol as described in: https://citeseq.files.wordpress.com/2019/02/cell_hashing_protocol_190213.pdf. Briefly, for the single-cell RNA-seq library of multi-time point FP-control MEFs, MEFs that were transduced with a pool of nine fluorescent proteins for different lengths of time (0 days - uninfected, 5 days, 15 days, and 21 days, respectively) were prepared separately into cell suspension. Additionally, MEFs 5 days after transduced with another three-fluorescent protein pool were also prepared into cell suspension. Cell hashing was performed on these five populations of cells as described previously (^74^). Briefly, cells were blocked with TruStain FcX™ (anti-mouse CD16/32) Antibody (Biolegend), and were then stained with TotalSeq™-A anti-mouse with Hashtag (Biolegend). Each of the five cell populations was stained with a different Hashtag. Cells were washed twice with PBS + 2% BSA + 0.01% Tween, before checked viability with TrypanBlue and counted. Based on the counts, the five populations were pooled in equal numbers, spun down, and resuspended in PBS + 0.04% BSA to reach the concentration of 1600 cells/µL. Finally, the pooled population was loaded for 10X Genomics scRNA-seq at a 2.3x overload (i.e., adding 10X recommended volumes of cells and water for a cell suspension of 700 cells/µL aiming for recovering 10 thousand cells, while in reality having a cell suspension of 1600 cells/µL).

To prepare gene expression as well as hashtag libraries for the control MEFs, we followed standard 10X Genomics protocol, with modifications to amplify the hashtags as described previously ^74^. Briefly, HTO PCR additive was spiked into the cDNA amplification PCR of the 10X protocol. During post-cDNA amplification PCR cleanup with SPRI beads (0.6x), whereas the supernatant after beads incubation was normally discarded, it was kept instead this time with another round of 2x SPRI cleanup to purify the hashtag PCR product. The purified product was amplified again with 2x KAPA Hifi PCR Master Mix, TruSeq DNA D7xx_s primer (containing i7 index) and SI-PCR primer (**Supplementary Table S8**). Finally, the PCR product was purified using a 1.6x SPRI cleanup and sequenced on a Nextseq sequencer.

#### Single cell RNA sequencing library preparation

14 days post-infection, infected MEFs were trypsinized and prepared for single cell RNA-seq following the standard 10X cell preparation protocol. The concentration of single cell suspension was adjusted to 500-1000 cells/μL and was loaded on the 10x Genomics Chromium system (10x Genomics, Pleasanton, CA) with the aim of generating 6000 to 10000 transcriptomes per channel (Chromium Single Cell 3′ Library & Gel Bead Kit v2, catalog number 120237). Single cell RNA sequencing libraries were constructed following the 10X Chromium Next GEM Single Cell 3’ Protocol with some modifications, which enables customized TF perturbation enrichment^75^. (**Supplementary Table S8**)

To prepare TF perturbation libraries: Aliquot 50-100 ng of purified cDNA amplification product (usually 2-4µl) from the 10X Genomics 3’ single-cell RNA-seq protocol. The cDNA is amplified for 13 cycles with primers (SI-PCR Primer from 10X Genomics 3’ single-cell RNA-seq protocol and Modified Nextera N7 Adapter) and 2x Kapa Hifi Mastermix (Roche #7958935001) (**Supplementary Table S8**). The PCR product is about 1 kb-long. It is purified with a 0.6x SPRIselect beads clean-up, following standard protocol, and sequenced on a Nextseq sequencer.

### COMPUTATIONAL METHODS

#### Reprogramming TF prediction

The raw fastq files from the Tabula Muris project were downloaded and reprocessed the same way as in-house datasets to minimize technical differences ^43^. Reprogramming TFs were predicted for every cell type as described in ^16^ with some modifications. In brief, entropy of each TF was calculated and used to access TF cell type specificity. In house primary datasets were used for cardiac cell type predictions ^16,76,77^. With the aim of sampling 10% of total TFs (∼200), we sought to maximize the number of cell types targeted. In the end, cardiomyocyte, lung endothelial cell, mesenchymal stem cell, neuroendocrine cell, skeletal muscle satellite cell, and stromal cell were chosen.

RSEM (version 1.3.3) was used to select the most abundant isoforms of the predicted reprogramming TFs in target primary cells.

#### Sequencing, basecalling and demultiplexing

Libraries were sequenced using the Illumina NextSeq 500/550 and NovaSeq 6000 (Illumina). The generated binary base call (BCL) files were demultiplexed and converted to standard FASTQ files using the mkfastq function from the Cell Ranger pipeline (version 5.0.1).

#### Read alignment and generation of gene expression matrix

Cell Ranger pipeline (version 5.0.1) was used to preprocess the 10X Genomics single-cell data with default parameters except the number of cells expected was set to 10,000 to generate the expression matrix. Scrublet was used to detect multiplets (version 0.2.3) ^78^.

Cell hashing demultiplexing

#### Perturbation detection

The detection and quantification of perturbations in single cells was performed using FBA ^79^ (version 0.0.12). The mismatch thresholds for cell and feature barcodes were set to 1 and 2, respectively. Demultiplexing was performed using the gaussian mixture model. Demultiplexing of barcoded antibody-labeled MEFs (cell hashing) results was performed using FBA.

#### Dimensionality reduction

Prior to normalization, non-informative cells and genes were filtered. Cells with less than 200 genes detected were excluded in the downstream analysis. For non-cardiac reprogramming assays, cells with more than 20% of UMIs from mitochondria were filtered and for cardiac reprogramming, the cutoff is 50%. Genes detected in fewer than 30 cells or with fewer than 60 UMIs in total across all single cells were filtered. Total UMI counts of single cells were normalized to the median UMI counts per cell and natural-logarithmized with a pseudocount of 1. The median-normalized log-transformed matrix was standardized (mean 0, standard deviation 1).

Principal component analysis (PCA) was performed using the Scanpy package (version 1.7.2) ^80^ with the median-normalized log-transformed and standardized matrix as input. Batch effect removal was performed using the harmony algorithm with default parameters (https://github.com/slowkow/harmonypy, version 0.0.5) ^81^. Each 10X Genomics RNA-seq library was treated as a different batch.

Uniform Manifold Approximation and Projection (UMAP) visualization was performed using UMAP with harmony-corrected PCA result as input (version 0.5.3, metric=“euclidean”, min_dist=0.1, spread=1.0) ^82^.

Minimum-distortion embedding (MDE) of perturbation cocktails was performed using PyMDE (version 0.1.18) ^83^. Expression matrix of perturbation cocktails was median-normalized and natural-logarithmized with a pseudocount of 1. Variable expression detection and regression out effects of total UMIs per cell was performed using Scanpy with default parameters. PCA was performed and used as input for pyMDE (n_neighbors=2, repulsive_fraction=5). The initial iterate was a 2-dimensional matrix generated by PHATE using PCA result (version 1.0.10) (knn=2, decay=40, n_landmark=2000, t=”auto”, gamma=1, mds_solver=“smacof”, knn_dist=“euclidean”, mds_dist=“euclidean”).

#### Clustering

Single cells were clustered using the leiden algorithm (leidenalg, version 0.9.1) implemented in the Scanpy package (version 1.7.2) ^80^. In brief, the harmony-corrected matrix was used to compute a neighborhood graph of observations (n_neighbours=30, method = “umap”, metric=“euclidean”). Then, the leiden algorithm was used to cluster single cells (resolution=None, flavor=“igraph”).

Hierarchical DBSCAN clustering of perturbation cocktails was performed using the hdbscan method implemented in dbscan with minPts setting to 10 (version 1.1.1) ^84^.

#### Differential expression analysis

### Binomial test

The binomial test based method used for testing if a gene is expressed more frequently between different groups of cells was described in Duan et al. 2019 ^16^ with small modifications.

### Pseudo-bulk

A pseudo-bulk approach was used to detect differential expression between two groups of single cells. Single cells belonging to the same groups were randomly combined (minimal cells: 30, maximum cells: 500). edgeR’ lrt method was used to detect differentially expressed genes between the pseudo-bulk replicates (version 3.38.4) ^85^.

#### Functional annotation and enrichment analysis

GO term enrichment was performed using Fisher’s exact test implemented in topGO (version 2.48.0; org.Mm.eg.db version 3.15.0). Reactome pathway enrichment was performed using ReactomePA (version 1.40.0; org.Mm.eg.db version 3.15.0; reactome.db version 1.81.0) ^86^.

#### Pseudotime construction

Monocle (version 2.24.1) was used for pseudotime analysis with differentially expressed genes between target primary cell types and control cells used for pseudotime ordering ^87^.

#### TF Interaction analysis

Genes detected in less than 30 cells or with less than 60 UMIs in total across all single cells were excluded for downstream analysis. UMI count matrix of perturbed cells were normalized to fluorescent protein infected control cells. For pairwise TF interactions, we required two-TF perturbed cells and their singlet constituents to have at least 10 cells. The median number of cells used for TF interaction interference is 595.Theil-Sen regression implemented in scikit-learn (version 1.2.0) was used for linear regression (max_subpopulation=1e5, n_subsamples=None, max_iter=1000, tol=0.001).

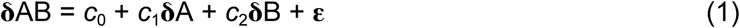

dcor package was used for distance correlation calculation (version 0.6). F-statistics was calculated to test the significance of regression coefficients in models. If the p-value associated with the F-statistics is less than 0.05, the interaction of the TF pair was characterized as follows:

##### Redundant

Variance Inflation Factors (VIF) was calculated to detect multicollinearity. If VIF is higher than 5, this pair is characterized as redundant.

##### Dominant

To determine the dominant effect of TF pairs (AB), the results of linear models (1, 2, 3) were taken into consideration.

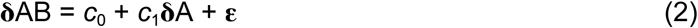

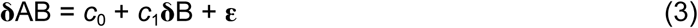

For a given TF pair (AB), if the *R*-Squared (*R*² or the coefficient of determination) of (2) is higher than that of (3) and *c*_1_ is higher than *c*_2_ in (1), then this TF Pair is characterized as A dominant; If the *R*² of (3) is higher than that of (2) and *c*_2_ is higher than *c*_1_ in (1), B is characterized as dominant.

##### Interactive

TF pairs (AB) were modeled using (4), and t-statistics of *c*_1_, *c*_2_ and *c*_3_ were calculated. If the p-value associated with the t-statistic of *c*_3_ is less than 0.05, this pair of TFs was characterized as interactive.

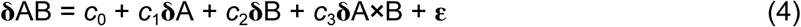

##### Neomorphic

Mean absolute error (MAE) from (1) was calculated for each TF pair. The largest 10% of TF pairs’ MAE were characterized as neomorphic.

#### Prediction

Prior to fitting classification models, low informative genes of single cells were filtered as described above. Then single cells were normalized and log-transformed. For each training (total 100 times), 20% of each group of single TF perturbed cells were randomly left out for validation. Two strategies were used. One is stochastic gradient descent classifier (SGDClassifier implemented in scikit-learn, version 1.3.1, LOSS=“log_loss”, PENALTY=“l2”), and the other is extreme gradient boosting (XGBoost, python implementation version 1.7.6, learning_rate=0.2, objective=“multi:softprob”, tree_method=“hist”).

## SUPPLEMENTAL TABLE LEGENDS

**Supplemental Table 1**: Transcription factor prioritization for each reprogramming assay.

**Supplemental Table 2**: Sequencing statistics for each reprogramming assay.

**Supplemental Table 3**: Sequencing statistics for TF combinations.

**Supplemental Table 4**: Differentially expressed genes in each perturbation cluster.

**Supplemental Table 5**: Reactome pathways and Gene Ontology terms used in Fig. 2G.

**Supplemental Table 6**: Statistics for linear models.

**Supplemental Table 7**: Summary of perturbation prediction accuracy.

**Supplemental Table 8**: Primers, TF ORFs, and constructs used in this study

## SUPPLEMENTAL FIGURE LEGENDS

**Supplemental Figure 1.**
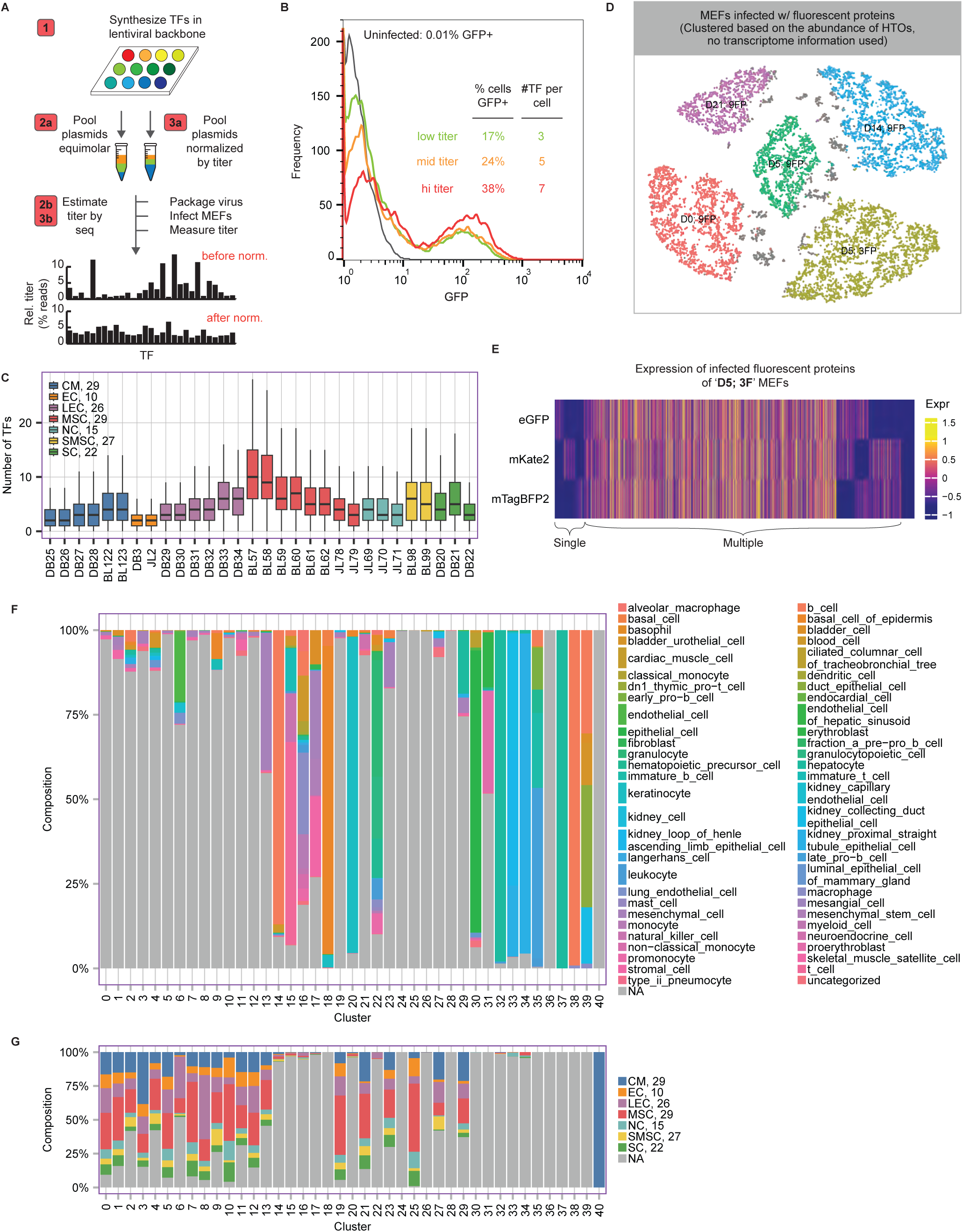
A. Schematic of iterative strategy to normalize lentiviral TF titers. As the lentiviral titer of each TF may be different, normalization of relative titers will be important for downstream analysis. To assess the relative titers of TFs in each reprogramming batch, we will create an equimolar pool of plasmids, package pooled lentivirus, infect MEFs for 7 days, and measure the titer for each TF by next-generation sequencing of TF barcodes from mRNA. We then adjust the concentration of each plasmid and iterate this analysis to confirm normalization of relative titers. B. Controlling perturbation complexity by controlling titer. We infected MEFs with a pool of 30 normalized TFs, quantified GFP+ cells by flow cytometry as a measure of absolute titer, and confirmed these measurements by scRNA-Seq. C. The median number of TFs in each batch of perturbation assay. D. Visualization of MEFs demultiplexing. Four batches of MEFs were labeled and pooled together prior to sequencing (cell hashing). The clustering of visualization was performed using the abundance of hashtag oligos. Colors indicate cell groups after demultiplexing. Grey dots indicate multiplets. E. Expression heatmap of infected fluorescent proteins of ‘D5; 3F’ MEFs (infected with 3 fluorescent proteins and cultured for 5 days). F. Cell type compositions of each cluster. Only cell type annotations from Tabula Muris were shown (non-grey colored). G. Cell type compositions of each cluster. Only cell type annotations from reprogrammed cells were shown (non-grey colored).

**Supplemental Figure 2.**
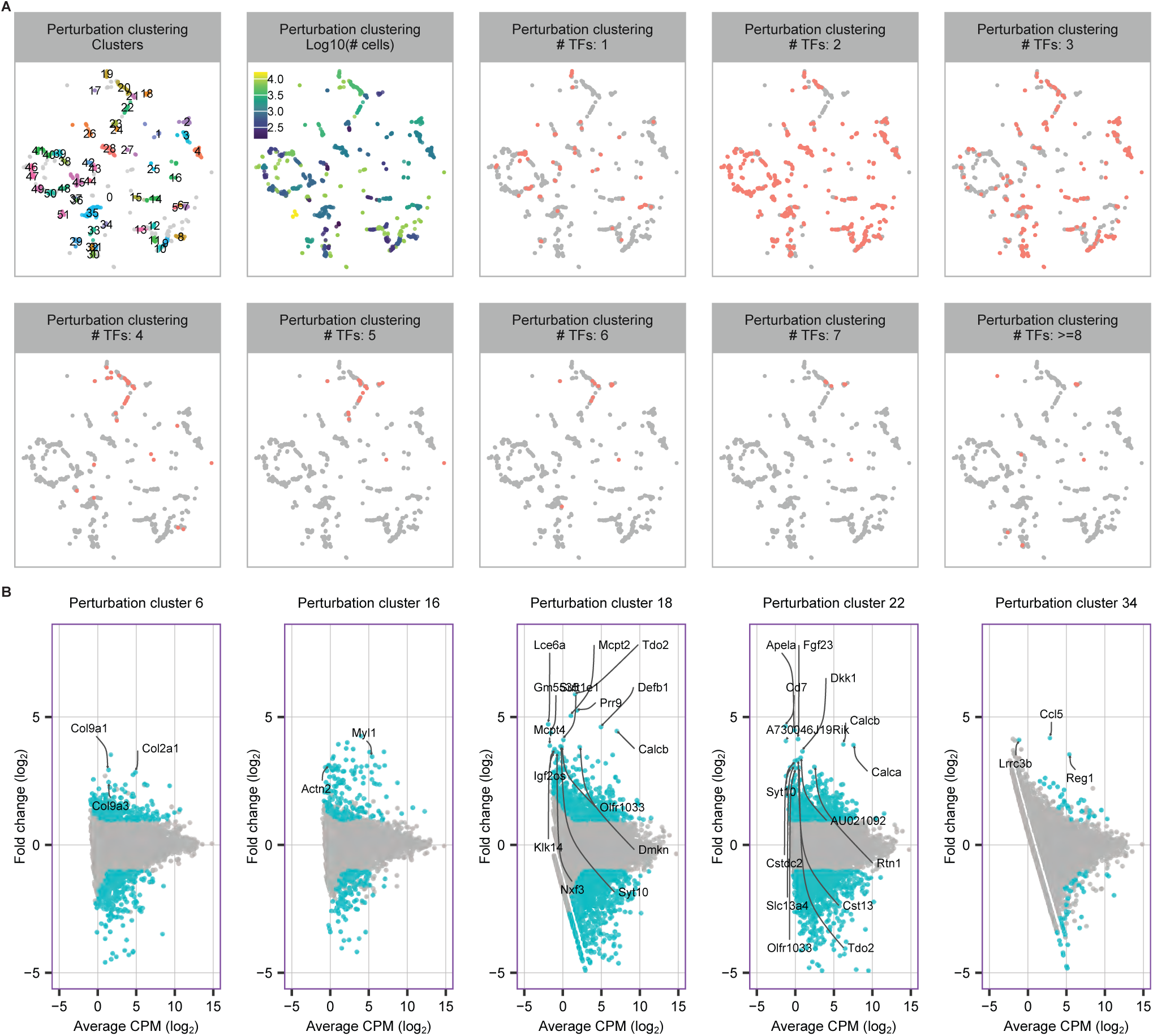
A. MDE visualization of perturbation clusters. From top left to bottom right are perturbation clusters, number of cells for each perturbation cluster, and number of TFs in each perturbation cluster. B. Differentially expressed genes (DEGs) for selected perturbation clusters. Teel indicates DEGs.

**Supplemental Figure 3.**
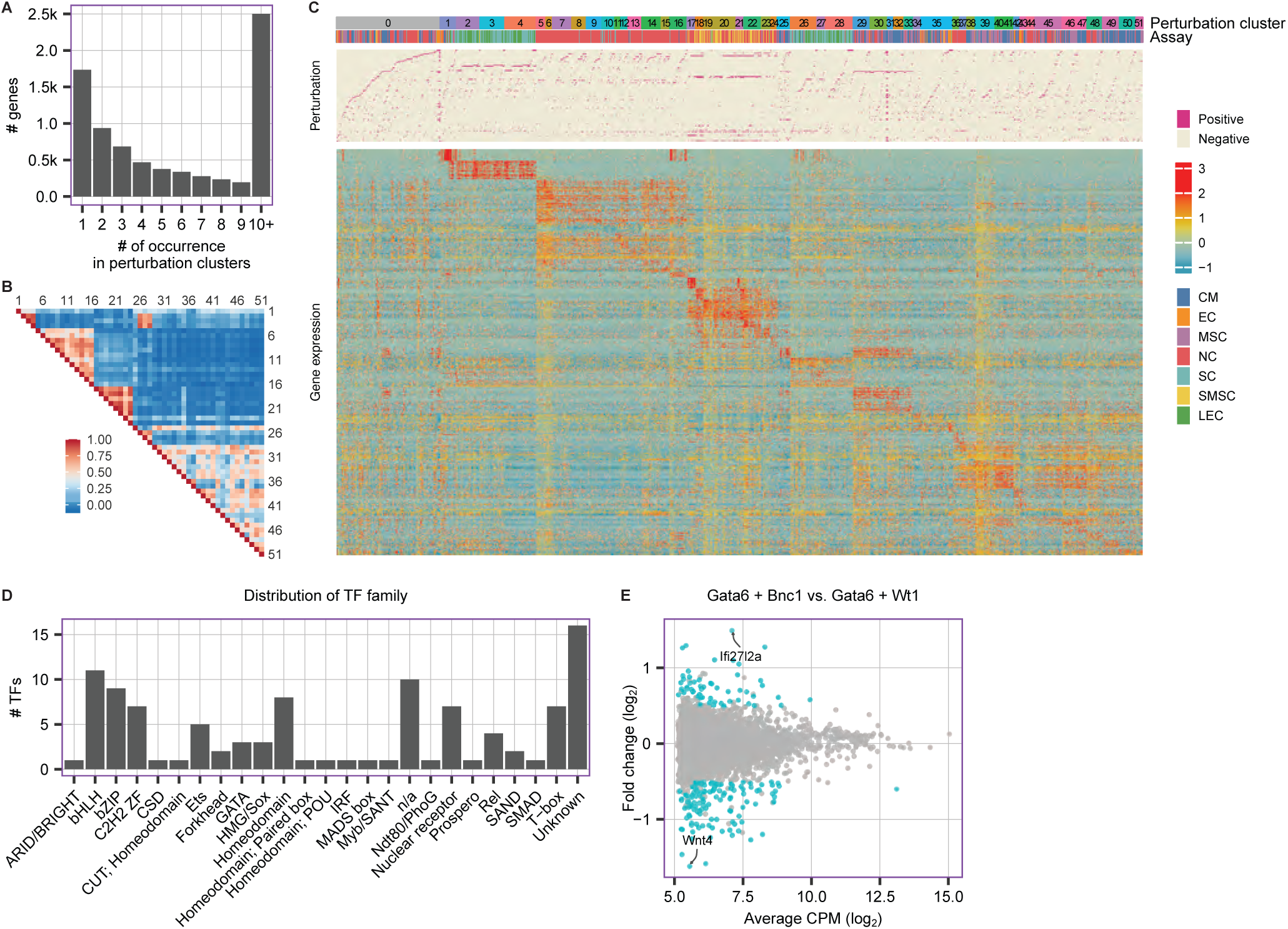
A. Across all differentially expressed genes identified in Fig. 3B, shown is the occurrence across modules). 1 indicates a gene is found only in one module. B. The heatmap indicates the correlation of TFs across perturbation clusters. C. Heatmap indicating gene expression patterns (bottom) across perturbation clusters. Each column indicates a distinct TF combination (middle), grouped by batch/assay and perturbation cluster (top). D. Distribution of the TF family annotations for the 105 TFs ^88^. E. The MA plot indicates the differentially expressed genes between TF pairs Gata6 + Bnc1 and Gata6 + Wt1 (teal). Several genes are indicated.

**Supplemental Figure 4.**
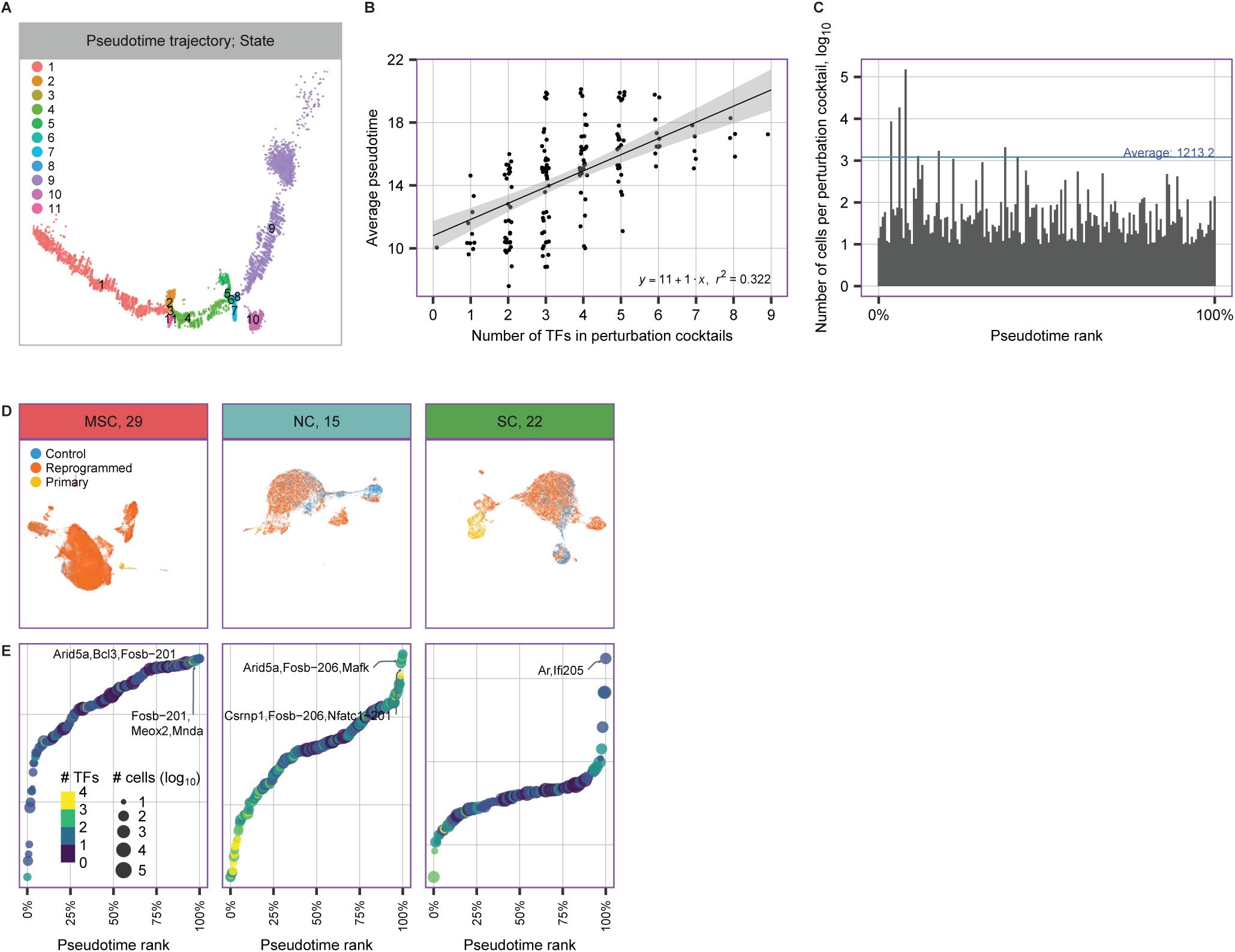
A. Pseudotime trajectory of epicardial reprogramming. Colors indicate different cellular states. B. The distribution of the numbers of TFs in perturbation cocktails and the average pseudotimes. C. The distribution of pseudotime ranking and the numbers of cells of these perturbation cocktails. D. Across all reprogramming batches and targeted primary cells, shown is the clustering of control MEFs (blue), MEF-derived cells (orange), and primary cells (yellow). E. Pseudotime analysis to rank putative TFs for cardiomyocytes, lung endothelial cells and mesenchymal stem cells reprogramming.

**Supplemental Figure 5.**
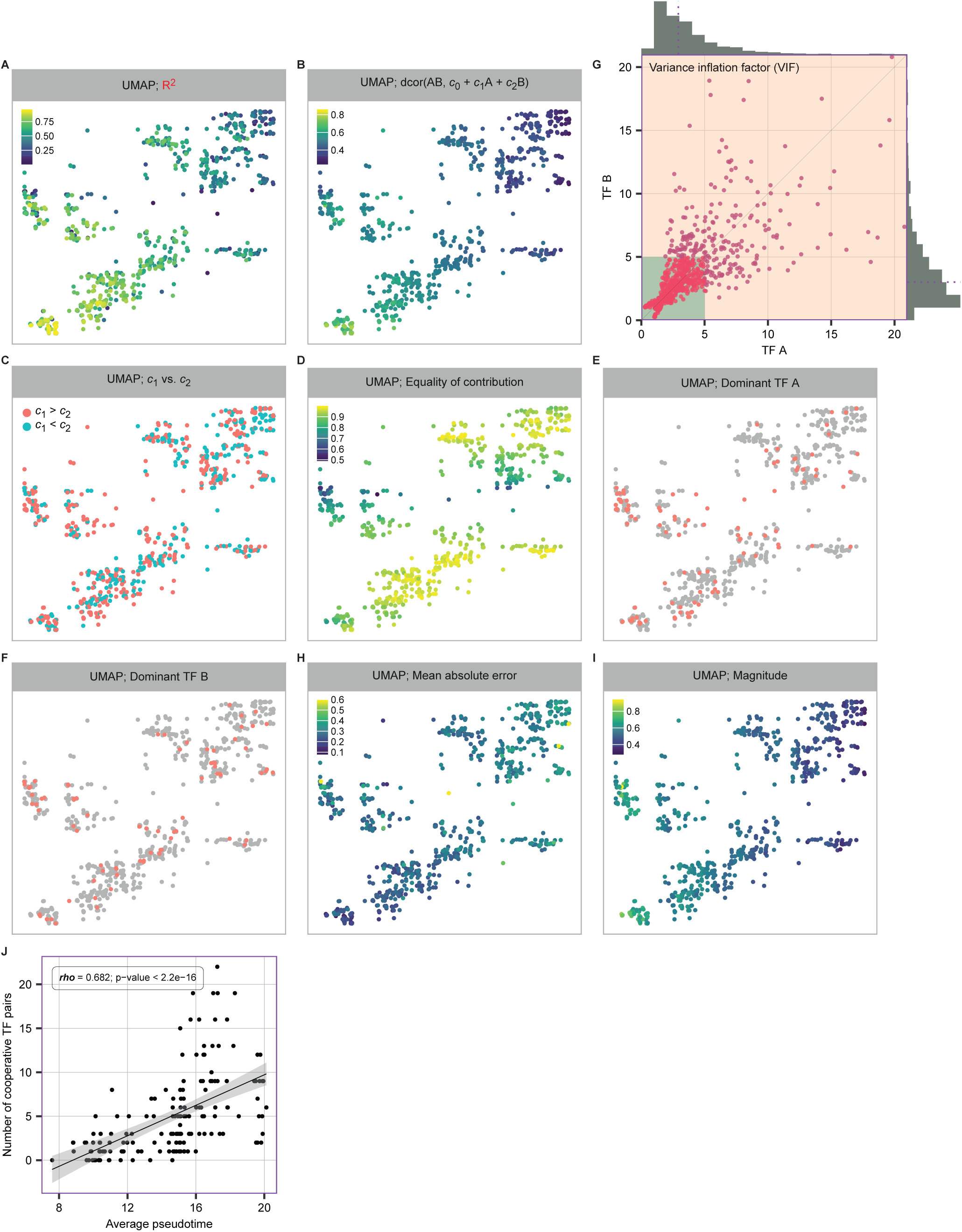
Visual embedding of TF pairs based on parameters of the linear model, the colors indicating R^2^ values (A), dcor correlation (B), relationship of *c*_1_ and *c*_2_ (C), equality of contribution (D), TF A dominant (E), TF B dominant (F), mean absolute error (H), magnitude (I) and variance inflation factor (G). J. The correlation of average pseudotimes and the number of cooperative TF paris in the perturbation cocktails.

**Supplemental Figure 6.**
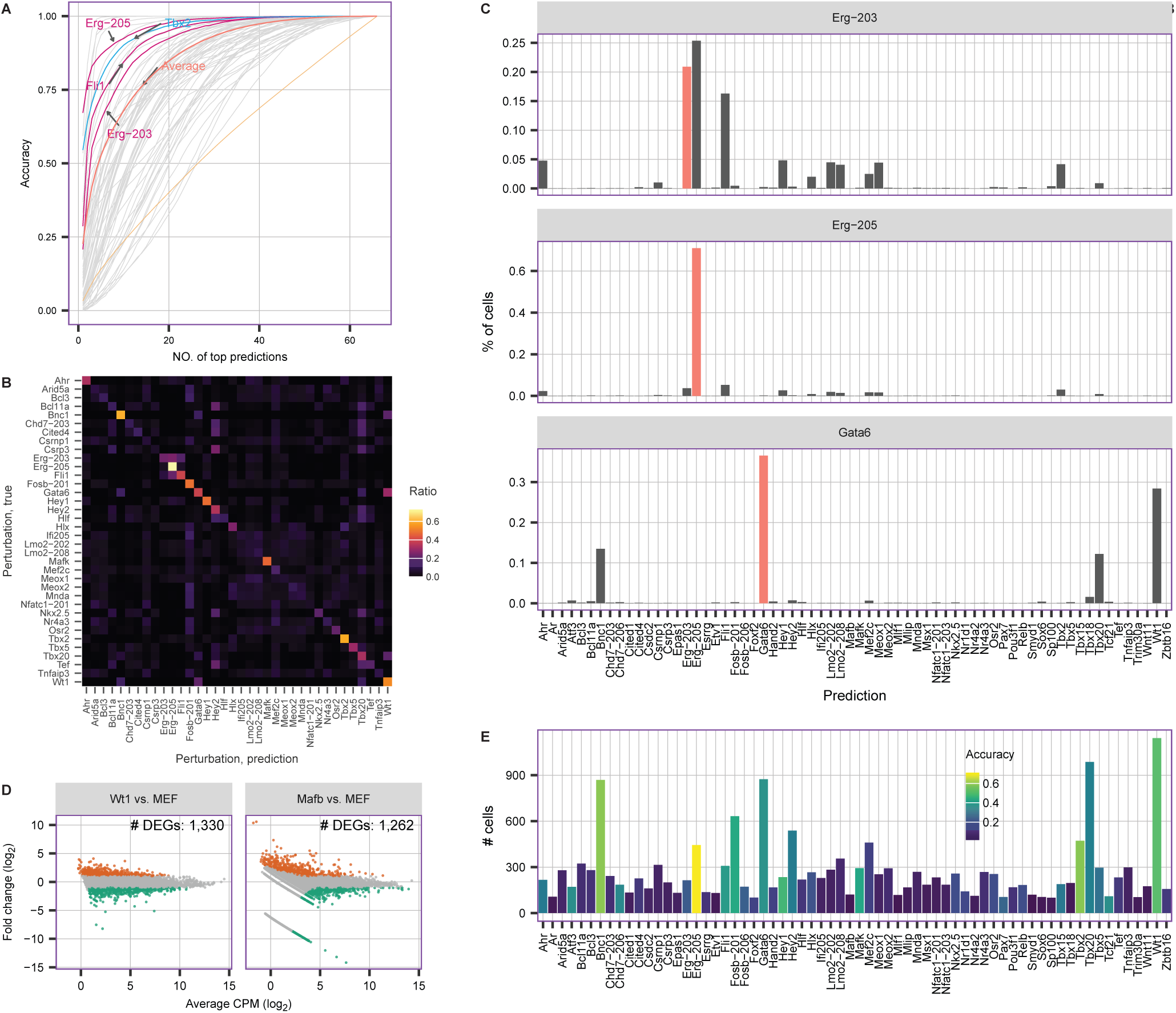
A. Prediction accuracy for 66 TFs using XGBoost algorithm with different cutoffs (0.2266). B. Heatmap visualization of the distribution of predictions (the sum of row is 1). C. The distributions of predictions for Erg-203, Erg-205, and Gata6. D. MA-plots of Wt1 and Mafb perturbed cells. Colors indicate differentially expressed genes. E. Number of cells for each single TF perturbation. Color indicates the prediction accuracy of each TFs.

